# The Alzheimer susceptibility gene *BIN1* induces isoform-dependent neurotoxicity through early endosome defects

**DOI:** 10.1101/2021.04.02.438184

**Authors:** Erwan Lambert, Orthis Saha, Bruna Soares Landeira, Ana Raquel Melo de Farias, Xavier Hermant, Arnaud Carrier, Alexandre Pelletier, Lindsay Davoine, Cloé Dupont, Philippe Amouyel, Amélie Bonnefond, Frank Lafont, Farida Adelfettah, Patrik Verstreken, Julien Chapuis, Nicolas Barois, Fabien Delahaye, Bart Dermaut, Jean-Charles Lambert, Marcos R. Costa, Pierre Dourlen

## Abstract

The *Bridging Integrator 1* (*BIN1*) gene is a major susceptibility gene for Alzheimer’s disease (AD). Deciphering its pathophysiological role is challenging due to its numerous isoforms. Here we observed in Drosophila that human BIN1 isoform1 (BIN1iso1) overexpression, contrary to BIN1iso8 and BIN1iso9, induced an accumulation of endosomal vesicles and neurodegeneration. Systematic search for endosome regulators able to prevent BIN1iso1-induced neurodegeneration indicated that a defect at the early endosome level is responsible for the neurodegeneration. In human induced neurons (hiNs) and cerebral organoids, *BIN1* knock-out resulted in the narrowing of early endosomes. This phenotype was rescued by BIN1iso1 but not BIN1iso9 expression. Finally, BIN1iso1 overexpression also led to an increase in the size of early endosomes and neurodegeneration in hiNs. Altogether, our data demonstrate that the AD susceptibility gene *BIN1*, and especially BIN1iso1, contributes to early-endosome size deregulation, which is an early pathophysiological hallmark of AD pathology.

## INTRODUCTION

Alzheimer’s disease (AD) is the most common form of dementia, characterized by two main cerebral lesions: the extracellular aggregation of the amyloid beta (Aβ) peptide into senile plaques and the intracellular aggregation of phosphorylated Tau into tangles. In addition, other cytopathological features specific to familial and sporadic AD can be also observed such as abnormally enlarged early endosomes in neurons (Cataldo et al., 2000). At the genetic level, familial AD is due to mutations in *APP, PSEN1* and *PSEN2*. Sporadic AD is a multifactorial disease exhibiting a strong genetic component with an estimated attributable risk of 60-80% (Gatz et al., 2006). For the last decade, our understanding of this genetic component has strongly progressed with the identification of 76 loci associated with the disease (Bellenguez et al., 2020). Among these loci, *BIN1* is the second susceptibility gene after *APOE* in terms of association (Bellenguez et al., 2020; Kunkle et al., 2019; Lambert et al., 2013).

*BIN1* encodes at least 20 exons subject to extensive differential splicing, generating multiple isoforms with different tissue distributions (Prokic et al., 2014). *BIN1* isoform1 (BIN1iso1) and *BIN1* isoform8 (BIN1iso8) are respectively expressed in the brain and skeletal muscles, the two tissues where *BIN1* is mostly expressed, whereas *BIN1* isoform9 (BIN1iso9) is ubiquitously expressed (GTEx portal, http://www.gtexportal.org). In the brain, BIN1iso1 and BIN1iso9 are the most abundant isoforms (Crotti et al., 2019; Taga et al., 2020). All *BIN1* isoforms possess the N terminal BIN1/Amphiphysin/Rvs (BAR) domain, involved in membrane curvature sensing and induction, the C terminal MBD domain for MYC-Binding Domain and the C-terminal SH3 domain, a protein-protein interaction domain, recognizing proline rich domain like the one in Tau (Prokic et al., 2014; Sottejeau et al., 2015). Muscle-specific isoforms contain a phosphoinositide-interacting (PI) domain, whereas brain-specific *BIN1* isoforms are mainly characterized by inclusion of exons encoding a CLAP domain involved in endocytosis and intracellular trafficking. In the brain, a complex expression pattern is also observed at the cellular level. *BIN1* expression is mainly observed in oligodendrocytes, microglial cells and neurons (Adams et al., 2016; Marques-Coelho et al., 2021; De Rossi et al., 2016). However, while neurons express high molecular weight isoforms including BIN1iso1, glial cells express lower molecular weight isoforms such as BIN1iso9 (De Rossi et al., 2016; Zhou et al., 2014).

AD-associated *BIN1* variants are non-coding and likely regulate *BIN1* expression (Chapuis et al., 2013). However, the deregulation of *BIN1* expression in the brain of AD cases is still highly debated. Some results indicate that overall *BIN1* expression is increased (Chapuis et al., 2013), or decreased (Glennon et al., 2013; Marques-Coelho et al., 2021), whereas more complex patterns have been also reported with a decrease in BIN1iso1 and a concomitant increase in BIN1iso9 expression (Holler et al., 2014). In addition, according to the pattern of expression, it is not clear if the observed variations of *BIN1* expression are a cause or a consequence of the neurodegenerative process. For example, the decrease in BIN1iso1 and increase in BIN1iso9 expressions may be a consequence of neuronal death and gliosis respectively as *BIN1* isoform variations correlated with neuronal and glial marker variations (De Rossi et al., 2016). Therefore, based on its global and/or isoform expression variation, it is difficult to assess whether BIN1 may be deleterious or protective in AD.

Importantly, impact of such global and/or isoform expression deregulations on the AD pathophysiological process has not yet been elucidated even if several hypothesis have been proposed: (i) modulation of Tau function and neurotoxicity though interaction of the BIN1 SH3 domain with the Tau proline-rich domain in a phosphorylation-dependent manner (Malki et al., 2017; Sartori et al., 2019; Sottejeau et al., 2015); (ii) modulation of Tau spreading through its role in endocytosis and intracellular trafficking or extracellular vesicles (Calafate et al., 2016; Crotti et al., 2019); (iii) regulation of the APP metabolism despite contradictory results in different models (Andrew et al., 2019; Ubelmann et al., 2017); (iv) regulation of synaptic transmission either in the presynaptic (De Rossi et al., 2020) or postsynaptic compartment (Schürmann et al., 2020).

Within this complex background, considering the numerous *BIN1* isoforms and the different functions regulated by this gene, it is, thus, pivotal to address isoform-specific functions of BIN1 towards a comprehensive understanding of its role in AD pathophysiology. For this purpose, we investigated the role of *BIN1* isoforms in neuronal cells by focusing on BIN1iso1, BIN1iso8 and BIN1iso9 in drosophila models and human induced neurons (hiNs) derived from induced pluripotent stem cells (iPSC).

## RESULTS

### Functional conservation of human BIN1 isoforms in Drosophila

In order to better understand the functional and pathological functions of BIN1 isoforms, we used the highly tractable Drosophila model which allowed easy generation and breeding of multiple transgenic models.

*BIN1* belongs to the amphiphysin family and in mammals, two genes compose this family: *amphiphysin I* (*AMPH*) and *amphiphysin II* (*AMPH2*) also named *BIN1*. Drosophila has only one ortholog called *Amphiphysin* (*Amph*) that is referred to as *dAmph* henceforth in this article. dAmph has 3 isoforms but only the BAR and SH3 domains are conserved in dAmph. Within this background, we generated 3 transgenic Drosophila lines expressing the human BIN1iso1, BIN1iso8 and BIN1iso9 isoforms. As a control, we also generated transgenic Drosophila lines expressing the longest dAmph isoform, dAmphA (Figure 1A). We obtained transgenic lines expressing identical basal level of BIN1 isoforms with two additional BIN1iso1 and BIN1iso9 lines expressing high BIN1 levels that we used for dose-dependent effects (Supplementary information, Figure S1).

**Figure 1.**
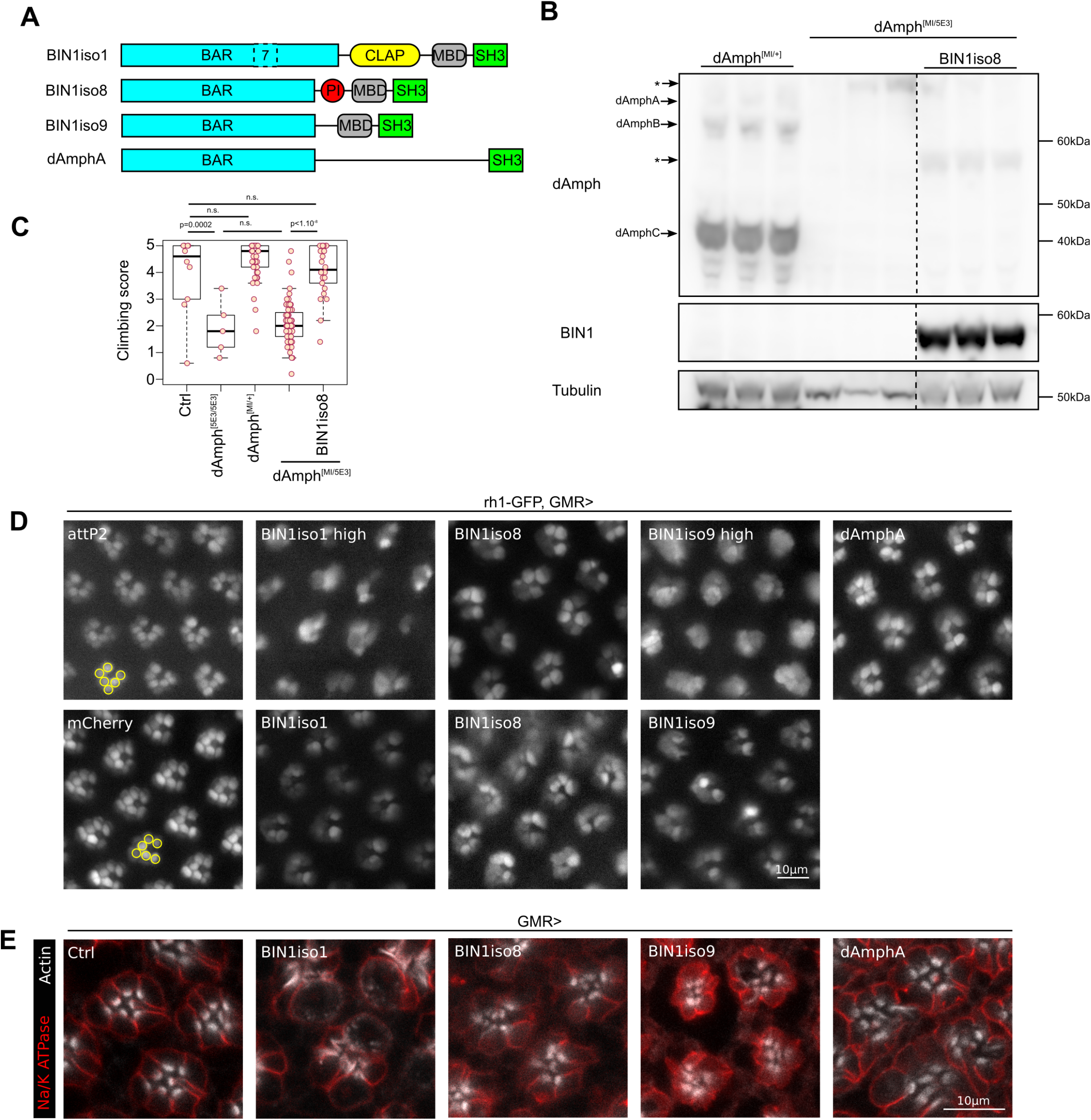
Functional conservation of human BIN1 isoforms in Drosophila. **(A)** Scheme of cerebral human BIN1 isoform1 (BIN1iso1), muscular human BIN1 isoform8 (BIN1iso8), ubiquituous human BIN1 isoform9 (BIN1iso9) and Drosophila BIN1, called Amphiphysin, isoformA (dAmphA) for which transgenic lines were generated. **(B)** Western blot analysis of Amph and BIN1 expression in dAmph^MI08903-TG4.0^/+, dAmph^MI08903-TG4.0/5E3^ and dAmph^MI08903-TG4.0/+^; UAS-BIN1iso8 fly thorax. dAmphA, dAmphB and dAmphC were expressed in the heterozygous dAmph^MI08903/+^ flies, whereas they were not detected in dAmph^MI08903/5E3^ flies (* background staining). BIN1iso8 was expressed in dAmph^MI08903-TG4.0/+^; UAS-BIN1iso8 flies. **(C)** Analysis of the climbing locomotor activity of 2 day-old flies with the indicated genotype. dAmph^MI08903-TG4.0/5E3^ flies exhibited a similar low climbing score as the null dAmph^5E3/5E3^flies. Expression BIN1iso8 rescued the locomotor defects of dAmph^MI08903-TG4.0/5E3^ flies (ANOVA p=2.37×10^−38^ with post-hoc Tukey, n.s. not significant). **(D)** Visualization of outer photoreceptor neuron rhabdomeres by cornea neutralization in 2-day-old flies expressing mCherry (as a control), BIN1iso1, BIN1iso8, BIN1iso9 and dAmphA under a GMR driver (attP2 is a control with an empty attP2 landing site and no UAS construct). While each ommatidium contained 6 outer photorceptors organized in a trapezoid shape (yellow circles) in the two control conditions, BIN1 isoforms and dAmphA expression resulted in a strong alteration in the number, shape and trapezoid organization of the rhabdomeres with a stronger effect for BIN1iso1 and BIN1iso9 high-expressing lines. **(E)** Immunofluorescence of whole-mount pupal retina expressing Luciferase (as a control), BIN1iso1, BIN1iso8, BIN1iso9 or dAmphA. They were labelled for the plasma membrane neuronal Na/K ATPase and F-actin. Contrary to the control, BIN1iso1, BIN1iso8, BIN1iso9 and dAmphA induced a strong accumulation of F-actin at the level of the rhabdomere.

We assessed the functional conservation of human BIN1 isoforms in Drosophila. Like human subjects harboring *BIN1* coding mutation and suffering myopathy (Nicot et al., 2007), *dAmph* null flies have locomotor defects, due to T-tubule morphogenesis defects in muscle cells (Leventis et al., 2001; Zelhof et al., 2001). We tested if the expression of muscle human BIN1iso8 could restore the locomotor performance of *dAmph* deficient adult transgenic flies assessed in the so-called climbing test. In addition to the null dAmph^5E3^ allele, we took advantage of a dAmph^MI08903-TG4.0^ allele that allows Gal4 expression under the control of *dAmph* endogenous promoter while stopping *dAmph* transcription (Supplementary information, Figure S2). We checked by western blot that *dAmph* expression was abolished in dAmph^MI08903-TG4.0/5E3^ compound heterozygous flies compared to dAmph^MI08903-TG4.0/+^ heterozygous flies and that BIN1iso8 could be expressed in this genetic background (Figure 1B). Then, we observed that dAmph^MI08903-TG4.0/5E3^ compound heterozygous flies had locomotor defects like dAmph^5E3/5E3^ null flies compared to control flies or dAmph^MI08903-TG4.0/+^ heterozygous flies (Figure 1C). Expression of human BIN1iso8 restored the locomotor abilities of dAmph^MI08903-TG4.0/5E3^ compound heterozygous flies to levels similar to the ones observed in control flies or dAmph^MI08903-TG4.0/+^ heterozygous flies (Figure 1C). Thus, human BIN1iso8 is able to rescue the locomotor functions of *dAmph* null flies indicating a functional conservation of human BIN1iso8 in Drosophila.

Next, we addressed the possible functional conservation of BIN1 isoforms in neuronal cells. Overexpression of dAmphA results in development defects of the light-sensitive photoreceptor neurons in Drosophila retina (Zelhof et al., 2001). These neurons possess a specialized compartment, called rhabdomere, which consists in an apical microvillar stack of actin-rich intricately folded membranes containing the light-sensing rhodopsin proteins. We tested if human BIN1 isoforms expression with the early eye-specific GMR driver could phenocopy the dAmphA-induced rhabdomere phenotype. The Drosophila eye is composed of 600-800 units called ommatida. Each ommatidium contains 6 outer photoreceptor neurons and 2 superimposed inner photoreceptor neurons. We expressed GFP in the outer photoreceptor neurons (Rh1 driver) and used the cornea neutralization technique to assess rhabdomere morphology (Dourlen et al., 2012). While we observed six rhabdomeres per ommatidium, of similar size and organized in a trapezoidal shape in the control condition, some rhabdomeres were missing and others were smaller or deformed in the dAmphA overexpressing condition (Figure 1D). Thus, we confirmed that expression of dAmphA alter rhabdomere morphogenesis. Expression of BIN1iso1, BIN1iso8 and BIN1iso9 resulted in a similar phenotype (Figure 1D). In addition, high levels of BIN1iso1 and BIN1iso9 exacerbated the phenotype indicating a dose-dependent effect. We also confirmed these results on whole-mount pupal retina dissection (Figure 1E). dAmphA-, BIN1iso1-, BIN1iso8- and BIN1iso9-overexpressing retina exhibited strong deformed accumulations of F-actin at the level of the rhabdomere. In conclusion, overexpression of all human BIN1 isoforms phenocopied dAmphA overexpression during the development of photoreceptor neurons suggesting functional conservation also at the neuronal level.

### Human BIN1iso1 is neurotoxic in Drosophila photoreceptor neurons

The neurotoxic effects of human BIN1 isoforms in the developing Drosophila prompted us to investigate whether a similar effect could also be observed after expression of BIN1 isoforms in adult Drosophila. To do so, we employed the outer photoreceptor-specific driver Rh1, active in photoreceptor neurons after rhabdomere morphogenesis at the end of pupal development. We quantified the number of outer photoreceptor neurons following cornea neutralization and rhabdomere visualization. We observed that young flies (1 day-old) had a normal number of outer photoreceptor neurons (6 per ommatidium) with normal morphology (Figure 2A, B). These observations indicated that the use of rh1 driver bypasses the developmental retinal defects observed using the GMR driver. We then observed that 4-week-old flies expressing BIN1iso1, but not the ones expressing BIN1iso8, BIN1iso9 and dAmphA, lost around half of their outer photoreceptor neurons (Figure 2A, B). This phenotype was not dose dependent as flies expressing basal or high BIN1iso1 levels had a similar outer photoreceptor neuron loss. Of note, flies expressing high levels of BIN1iso9 had nearly no loss of outer photoreceptor neurons (Figure 2 A-D). In conclusion, expression of BIN1iso1 induced a progressive neurodegeneration in adult Drosophila photoreceptor neurons and this effect was isoform-specific and not dose-dependent.

**Figure 2.**
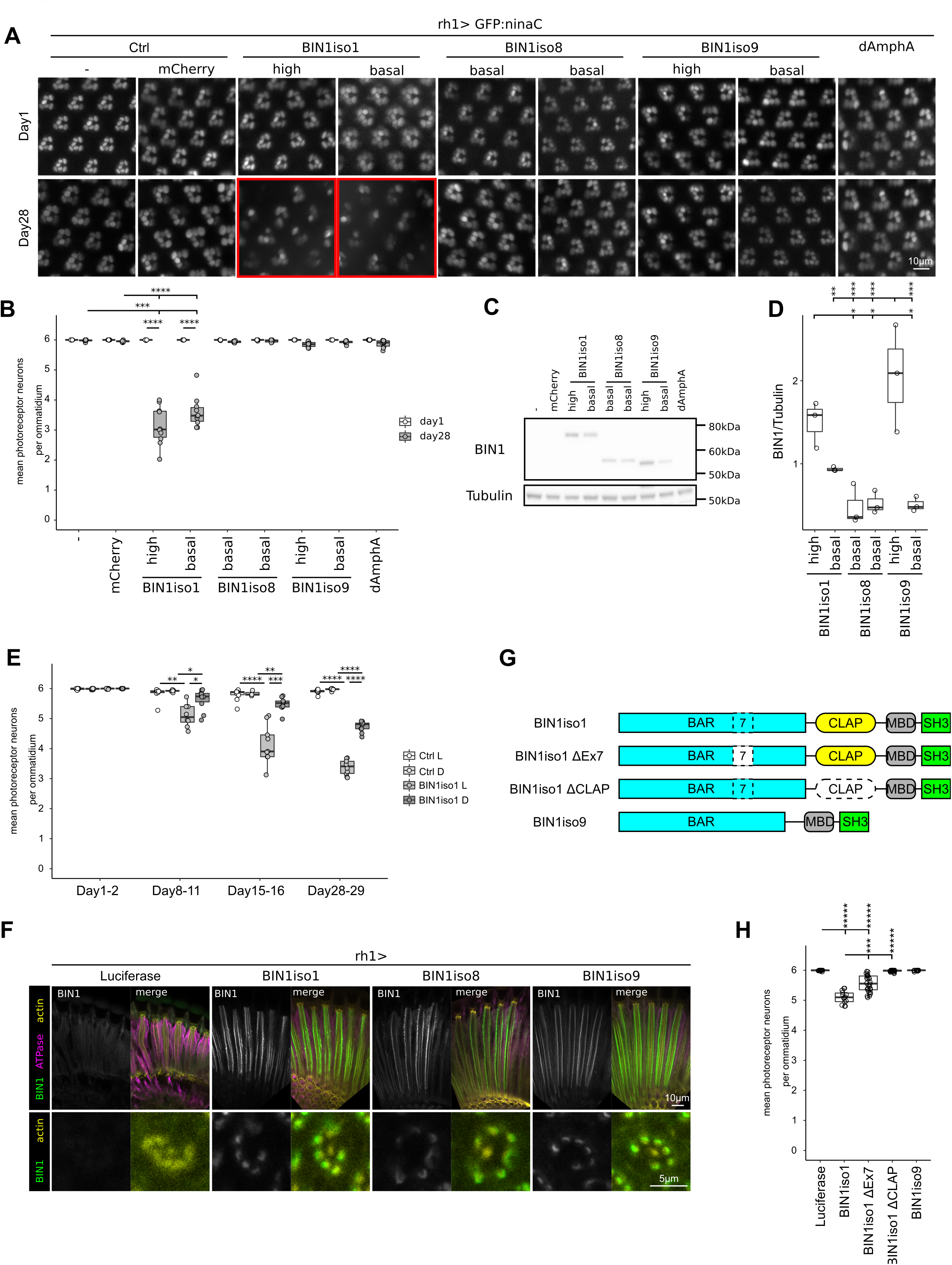
Human BIN1iso1 is neurotoxic in Drosophila photoreceptor neurons. **(A)** Visualization of retina photoreceptor neurons expressing BIN1 isoforms (rh1 promoter) by cornea neutralization in living flies. **(B)** Quantification (Kruskal Wallis p=0.013 followed by Mann-Whitney comparison, *** p<0.001, **** p<0.0001). Contrary to BIN1iso8 and BIN1iso9, BIN1iso1 expression induced a progressive age-dependent neurodegeneration, which was not dose-dependent. **(C-D)** Western blot analysis of BIN1 isoforms expression in the retina and quantification (ANOVA p= 0.00017, with post-hoc Tukey, * p<0.05, ** p<0.01, *** p<0.001). **(E)** Quantification of BIN1iso1-induced photoreceptor neuron degeneration over 4 weeks under a 12h/12h light/dark cycle (Ctrl L and BIN1iso1 L) or under constant darkness (Ctrl D and BIN1iso1 D) (Kruskal Wallis p=0.0005344 for Day8-11, p=3.264×10^−05^ for Day15-16, p= 1.36×10^−07^ for Day28-29, followed by Mann Whitney comparison, * p<0.05, ** p<0.01, *** p<0.001, **** p<0.0001). Loss of light did not prevent neurodegeneration although the intensity of degeneration was reduced. (**F**) Localization of BIN1 isoforms in one-day-old fly photoreceptor neurons. Luciferase was used as a control. Na/K ATPase staining labelled the plasma membrane and actin staining mostly labelled rhabdomere of photoreceptor neurons. Upper panels are longitudinal views of retina, whereas lower panels exhibit sectional view of ommatidium. **(G-H)** Scheme of the truncated BIN1Iso1 tested protein and quantification of their toxicity in 15 day-old flies (Kruskal Wallis p=5.686×10^−11^ followed by Mann Whitney comparison, *** p<0.001, ***** p<0.00001). Loss of the CLAP domain totally abrogated BIN1iso1 toxicity.

Since photoreceptor neurons are specialized neurons highly dependent on the phototransduction cascade (Wang and Montell, 2007), we wondered if a defect in this phototransduction cascade may be the cause of the neurodegeneration. For this purpose, we assessed whether the degeneration was light-dependent. We let BIN1iso1-expressing flies get old in the normal 12h/12h light/dark cycle or under constant dark condition for 4 weeks. Light absence did not prevent outer photoreceptor neurodegeneration even if occurring at a lesser extent (Figure 2E). This indicated that the phototransduction cascade was not the main cause of the neurodegeneration and that increased global neuronal activity favors BIN1iso1 induced neurodegeneration.

We asked why BIN1iso1 was more toxic for photoreceptor neurons than the other BIN1 isoforms. We wondered whether it could originate from the subcellular localization of BIN1 isoforms. We labelled retinas of one-day old flies expressing either BIN1iso1, BIN1iso8 or BIN1iso9 for BIN1 (Figure 2F). All BIN1 isoforms were similarly enriched at the base of photoreceptor neuron rhabdomeres. We next wondered what in the sequence of BIN1iso1 makes it neurotoxic. BIN1iso1 differs from BIN1iso9 by two sequences: (i) Exon7 in the BAR domain and (ii) the neuronal specific CLAP domain. We tested the neurotoxicity of truncated BIN1iso1 forms for these two sequences (Figure 2G) after generating the corresponding transgenic flies (Figure S3) and observed that loss of the CLAP domain abrogated outer photoreceptor neurodegeneration contrary to Exon7, which only partially rescued photoreceptor neurons (Figure 2H). Hence the BIN1iso1-induced degeneration depends on its CLAP domain and to a lesser extent on the Exon7 in the BAR domain. Since the CLAP domain is known to interact with AP-1 adaptin, AP-2 adaptin and the Clathrin Heavy Chain, which are involved in endocytosis and intracellular trafficking (Huser et al., 2013; Ramjaun and McPherson, 1998; Wigge et al., 1997), this suggested that the cause of the degeneration could be a defect in endocytosis/intracellular trafficking.

### BIN1iso1 induces vesicle accumulation in photoreceptor neurons

To further understand the cause of the BIN1iso1-induced degeneration, we analyzed the degenerating photoreceptor neurons by electron microscopy. While neurons in 15-day-old flies either expressing luciferase, BIN1iso9 or dAmphA did not show any abnormalities, the degenerating neurons in BIN1iso1 flies exhibited a strong accumulation of vesicles (Figure 3A, Figure S4A). These vesicles were of various sizes, some of them nearly as big as a complete ommatidium. The cytoplasm of some photoreceptor neurons was filled with vesicles, compacting it against the plasma membrane, which seemed intact. The vesicles were surrounded by a single membrane and not a double membrane as observed in autophagosome (Figure 3A) and did not contain any specific content. Rhabdomeres of BIN1iso1 photoreceptor neurons were disintegrated, whereas the chromatin seemed normal although the nucleus was frequently squeezed between vesicles (Figure 3A, Figure S4B). Eventually, photoreceptor neurons died (Figure S4D). Hence, ultrastructural analysis of degenerating photoreceptor neurons indicated that the neurodegeneration induced by BIN1iso1 is characterized by a strong accumulation of single membrane vesicles of unknown origin.

**Figure 3.**
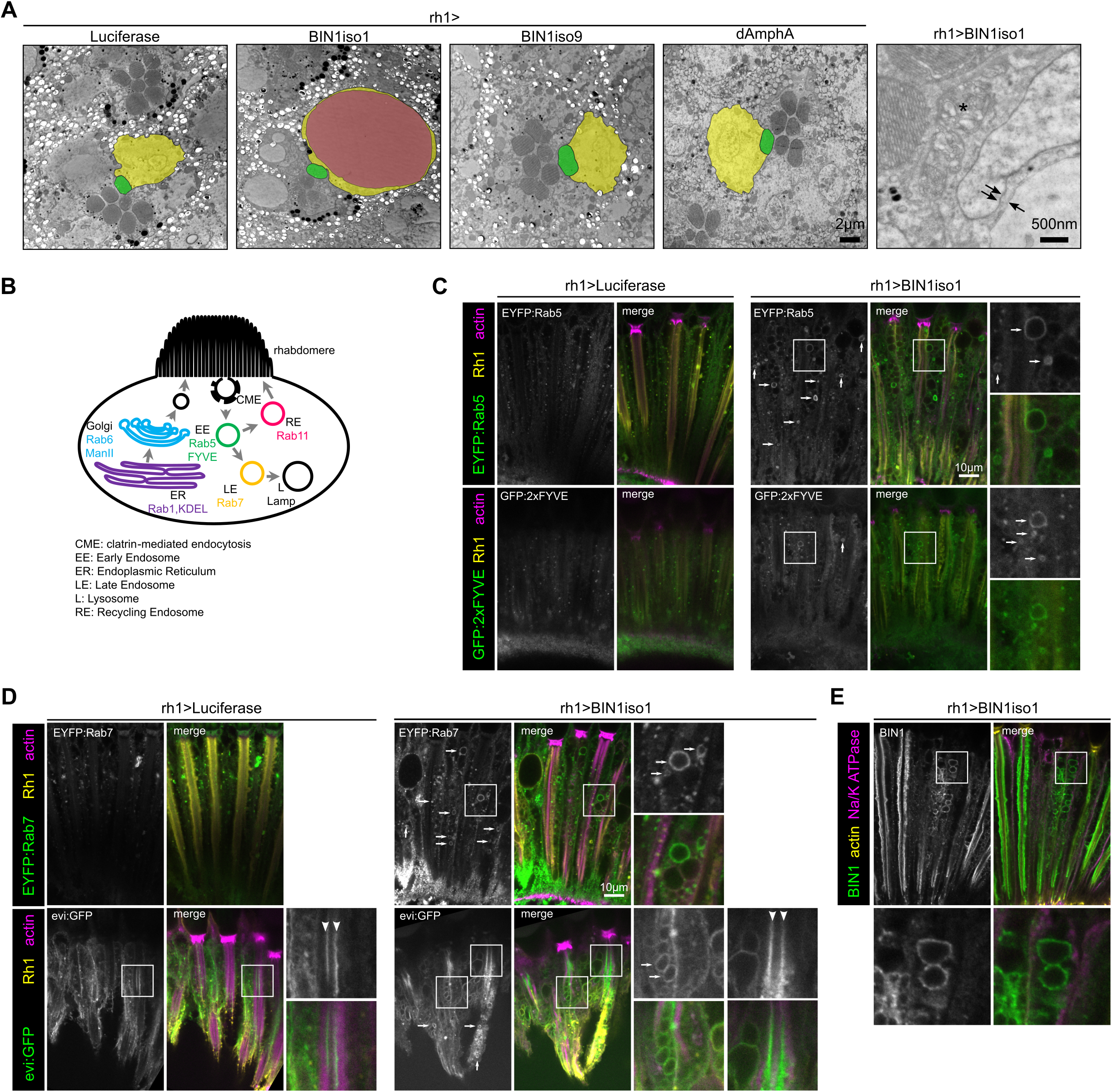
BIN1iso1-induced degeneration is characterized by a strong accumulation of vesicles showing endosomal markers. **(A)** Electron microscopy images of retina expressing Luciferase (Ctrl), AmphA, BIN1iso1 and BIN1iso9 from 15-day-old flies. 1 to 2 ommatidia transversal sections are seen on the 4 left images. One photoreceptor neuron is outlined per image, its cytoplasm and nucleus being highlighted in yellow and its rhabdomere in green. Each ommatidium contains 6 outer photoreceptor neurons whose rhabdomeres are organized in a trapezoidal shape, with a 7th rhabdomere in the middle corresponding to one of the two inner photoreceptor neurons. In the BIN1iso1 condition, only 4 rhabdomeres can be seen. In addition, the cytoplasm of the photoreceptor is filled with vesicles, some of them (highlighted in magenta) are very big. These vesicles are surrounded by a single membrane (arrow on the right panel) as compared to the nuclear envelope (double arrow). Their rhabdomere is usually disintegrating (*). **(B)** Scheme of the organelles tested in photoreceptor neurons with the markers used. **(C-D)** Images of photoreceptor neurons expressing BIN1iso1 and EYFP-tagged endogenous Rab5 and GFP:2xFYVE, as early endosome markers **(C)**, or EYFP-tagged endogenous Rab7 and evi:GFP, as late endosome/multivesicular body markers **(D)**. Rh1 and actin staining, respectively yellow and magenta in the merge image, are used to label retina structure of 15-day-old flies. The photoreceptor neurons expressed Luciferase as a control or BIN1iso1. In the BIN1iso1 conditions, many vesicles were positive for the tested organelle markers (arrows, green staining in merge images). The extracellular inter-rhabdomeric staining for the evi:GFP marker (arrowhead) corresponds to exosomes. (**E**) Images of photoreceptor neurons expressing BIN1iso1 stained for BIN1, the photoreceptor neuron plasma membrane Na/K ATPase and actin in 7-day-old flies. BIN1iso1 is localized at the base of the rhabdomeres and around some abnormal vesicles as seen in the inset.

We next evaluated the nature of these vesicles by immunofluorescence using specific organelle GFP- or YFP-tagged markers for endoplasmic reticulum (ER), Golgi, plasma membrane, early endosome (EE), late endosome/multivesicular body (LE/MVB), recycling endosome (RE), lysosome and autophagosome (Dunst et al., 2015) (Figure 3B, Figure S5). In 15-day-old flies, many BIN1iso1-induced vesicles were positive for EYFP:Rab5 and GFP:2xFYVE, markers of early endosome and for EYFP:Rab7 and evi:GFP, markers of late endosome/multivesicular body (Figure 3C, D). These different markers labelled small to middle size vesicles at the exception of evi:GFP which tended to label bigger vesicles. Some big vesicles were also exceptionally labelled by the lysosomal Lamp2:GFP marker and corresponded to rare multilamellar bodies observed by electron microscopy (Figure S4C, S5G). Of note, we also noticed that the extracellular space between rhabdomeres of the 8 photoreceptor neurons in the middle of ommatidia, called the interrhabdomeric space, was positive for evi:GFP in both the control and BIN1iso1 conditions (arrowhead Figure 3D). Since evi:GFP labels exosomes either within the multivesicular bodies or in the extracellular environment (Bartscherer et al., 2006; Lauwers et al., 2018), evi:GFP interrhabdomeric staining likely corresponded to released exosome which did not appear to be compromised by BIN1iso1 expression. In addition, we observed staining for BIN1 around some vesicles suggesting a potential direct effect of BIN1iso1 on the membrane dynamics of these vesicles (Figure 3E). In conclusion, these results suggested that BIN1iso1-induced vesicles accumulation originated from a blockade at the level of early endosome and/or late endosome.

### BIN1iso1 induces neurodegeneration through blockade of the early endosome trafficking in drosophila photoreceptor neurons

To test if the intracellular trafficking defects were responsible for the neurodegeneration phenotype, we tested if regulators of the endosome trafficking could rescue BIN1iso1-induced neurodegeneration. We tested regulator of early endosome (Rab5), recycling endosome (Rab11), late endosome (Rab7 and Rab9) and lysosome (subunits of the V-ATPase) (mostly a collection of UAS transgenes expressing wild-type, constitutively active and dominant negative Rab proteins (Zhang et al., 2007)). We observed an inhibition of BIN1iso1-induced neurotoxicity through a rescue of photoreceptor neurons for regulators of the early endosome Rab5 and recycling endosome Rab11 (Figure 4A, C, D). Modulation of the late endosome regulators Rab7 and Rab9, and of lysosome V-ATPase did not modify BIN1iso1-induced neuronal loss (Figure 4E-F). Of note, we checked by western blot that the rescue effect of Rab5 and Rab11 was not due to a decrease in BIN1iso1 expression, consecutive to a dilution of the Gal4 between the multiple UAS constructs (Figure 4B). We further tested the constitutively active (CA) and dominant negative (DN) forms of Rab5 and Rab11, respectively named Rab5^CA^, Rab5^DN^, Rab11^CA^ and Rab11^DN^. Surprisingly Rab5^DN^ rescued BIN1iso1-expressing photoreceptor neurons, whereas Rab5^CA^ had no effect (Figure 4C). This indicated that, although counterintuitive, overexpression of wild-type Rab5 resulted in a loss of function of Rab5 and that loss of function of Rab5 is protective against BIN1iso1-induced neurodegeneration. We confirmed this result when removing one copy of Rab5 (Rab5^2/+^) or knocking down Rab5 (Rab5^HMS00145^) (Figure 4C). Contrary to Rab5, Rab11^DN^ increased BIN1iso1-induced neuronal loss (although not significant) (Figure 4D). A gain of function of Rab11 seemed, therefore, protective against BIN1iso1-induced neurodegeneration. Altogether, because the neurodegeneration was rescued by modulation of regulators of early and recycling endosomes, these results indicated that BIN1iso1-induced photoreceptor neuron degeneration is due to a defect in the early endosome trafficking.

**Figure 4.**
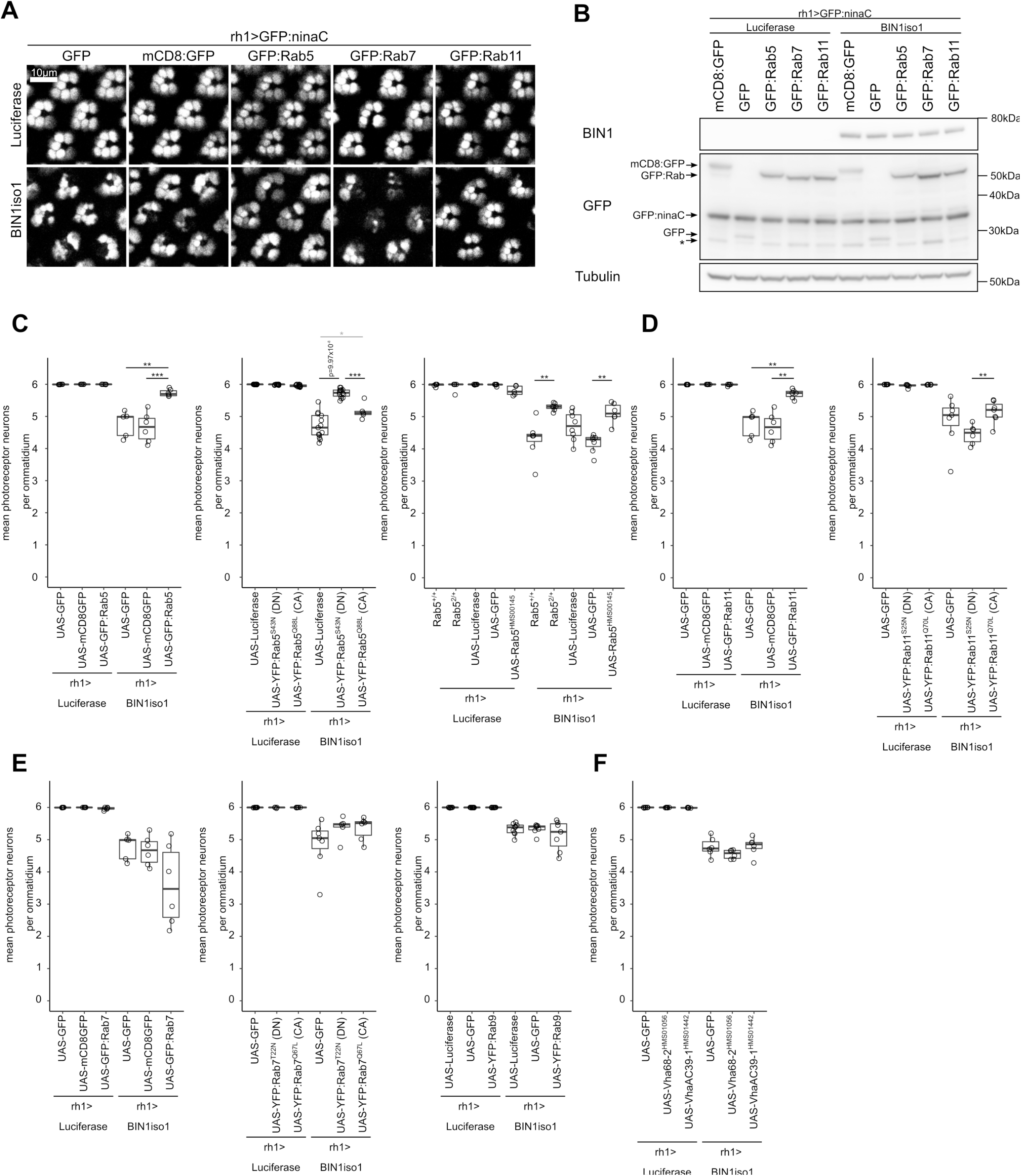
BIN1iso1-induced neurodegeneration is rescued by regulators of the intracellular trafficking. **(A)** Representative images of 15-day-old retina expressing BIN1-1 or luciferase (as a control) and constructs modulating Rab5, Rab7 and Rab11 activities. **(B)** Western blot analysis of BIN1, GFP derivatives in the corresponding conditions showing that BIN1 is not decreased in the conditions in which photoreceptor neurons are rescued. * non-specific band. **(C-F)** Quantification of the BIN1-1-induced neurodegeneration upon respective modulation of the early endosome regulator Rab5 activity, the recycling endosome regulator Rab11 activity, the late endosome regulator Rab7 and Rab9 activities and the lysosomal ATPase activity. Statistical analysis was performed using a Kruskal Wallis test (p=0.003643 for Rab5, p=6.169×10^−06^ for Rab5^DN^ and Rab5^CA^, p=0.0001721 for Rab5 mutant and knockdown, p=0.003643 for Rab11, p=0.03487 for Rab11^DN^ and Rab11^CA^, p=0.1408 for Rab7, p=0.24 for Rab7^DN^ and Rab7^CA^, p=0.8959 for Rab9, p=0.1394 for lysosomal ATPase subunit knockdown) followed by Mann Whitney comparison (* p<0.05, ** p<0.01, *** p<0.001).

### Generation and characterization of *BIN1 WT* and *KO* human induced neurons

Next, we wondered whether the role of BIN1iso1 in endosome trafficking was conserved in Human. To address this possibility, we generated human isogenic *BIN1* wild type (WT) and knockout (KO) induced pluripotent stem cells (iPSCs) lines by CRISPR/Cas9 technology. Both iPSC lines showed similar expression of pluripotency cell markers (SOX2 and SSEA4, Figure 5C) and growth rates (Figure 5D). *BIN1* WT iPSC expressed low molecular weight BIN1 isoforms, including likely BIN1iso9 (Figure 5I). We then employed these cells to generate iPSC-derived neurons both in bi-dimensional (2D) cell cultures and in cerebral organoids.

**Figure 5.**
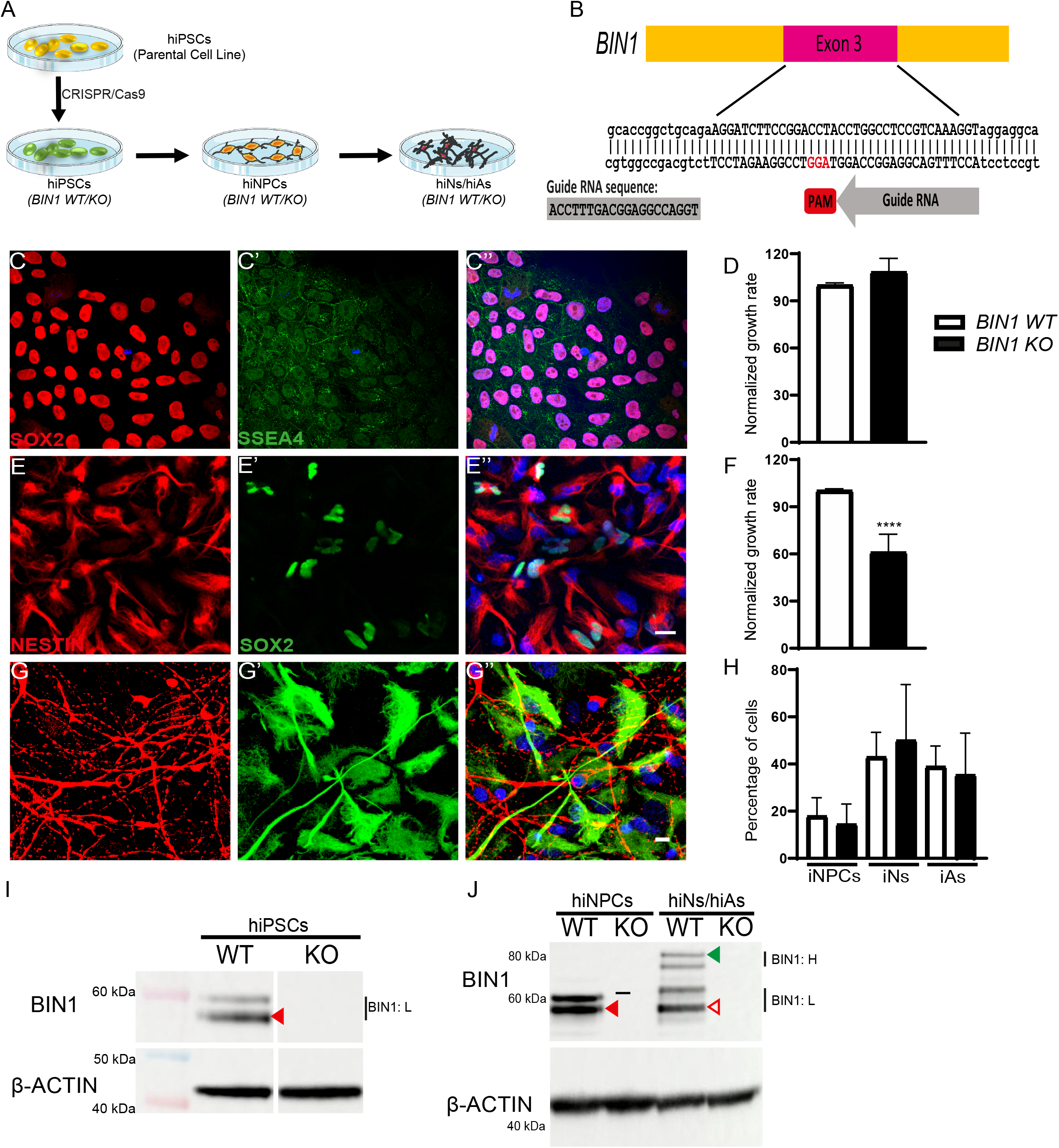
Characterization of hiPSCs and their cell derivatives. **(A)** Schematic showing the production of *BIN1* WT and KO iPSCs from parental cell line using CRISPR/Cas9 technology. These iPSCs, in turn, were used to generate intermediate iNPCs, and subsequently, mixed cultures of iNs and iAs. **(B)** Representation of exon 3 region of BIN1 was targeted for the production of *BIN1* WT and KO iPSCs by CRISPR/Cas9 technology. The guide RNA sequence is shown (grey). **(C-C’’)** Representative images showing pluripotency markers SOX2 (red) and SSEA4 (green), and stained with DAPI (C”) in *BIN1* WT and KO iPSCs. **(D)** Plot showing the cell proliferation numbers between *BIN1* WT and KO iPSCs (N= 4 independent cell passages; p=0.77, Unpaired t-test). **(E-E’’)** Representative images showing NPC markers NESTIN (red) and SOX2 (green), and stained with DAPI (E”, Scale Bar = 20 µm) in *BIN1* WT and KO iNPCs. **(F)** Plot showing the cell proliferation numbers between *BIN1* WT and KO iNPCs (N= 9 independent cell passages: ****p<0.0001, Unpaired t-test). **(G-G’’)** Representative images showing neuronal marker MAP2 (red), astrocytic marker GFAP (green) and stained with DAPI (G”, Scale Bar = 10µm) in 6-week-old *BIN1* WT and KO iNs and iAs. **(H)** Plot showing the percentage of cells in different cell populations – iNPCs, iNs, and iAs. (ANOVA _F(5,12)_, p=0.45). **(I)** Immunoblot validating the genotype of iPSCs for WT and KO. Band indicated on the blot for the WT cells (red arrowhead) indicates BIN1-light (BIN1:L) isoforms of BIN1. **(J)** Immunoblot showing the expression ofBIN1:L isoforms (solid red arrowhead) in WT iNPCsand both BIN1:L (hollow red arrowhead) and BIN1-heavy (BIN1:H; solid green arowhead) isoforms in 6-week old mixed cultures of iNs and iAs.

Following neural induction, both *BIN1* WT and KO human induced neural progenitor cells (iNPCs) expressed similar levels of NESTIN and SOX2 (Figure 5E). Although *BIN1* KO iNPCs showed a significant reduction in growth rate when compared to WT (Figure 5F), both iNPCs lines were expanded up to 10 passages and readily generated similar proportions of induced neurons (iNs) and astrocytes (iAs) upon differentiation conditions (Figure 5G-H). WT iNPCs mainly expressed low molecular weight BIN1 isoforms, whereas in differentiated cultures both low and high molecular weight isoforms, probably corresponding to BIN1iso1 and BIN1iso9, were expressed (Figure 5J). The expression pattern observed for BIN1 isoforms likely reflects the mixed composition of the differentiated cell cultures at 6 weeks, comprising both neurons and astrocytes, which mainly express BIN1iso1 and BIN1iso9, respectively (Zhou et al., 2014).

Likewise, *BIN1* WT and KO iPSCs were able to generate cerebral organoids with no obvious difference in size and composition (Figure 6A-B). Expression of BIN1 protein in 190-day-old *BIN1* WT cerebral organoids was confirmed by western blot and showed a similar pattern as the one described in 2D cell cultures (Figure 6C). Next, using single-nucleus RNA-sequencing (snRNA-seq), we observed that cerebral organoids of both genotypes contain all major neural cell types, with no significant difference in the proportions of cell types (Figure 6D-E; Figure S6). We then performed differential gene expression analysis for all different cell types identified in *BIN1* WT and KO cerebral organoids using DESeq2 (Love et al., 2014). We observed 41 differentially expressed genes (DEGs; 0.7<FC>1.3 and FDR<0.01) in glutamatergic neurons and 43 DEGs in astrocytes (Figure 6G-H and Table S1). All the other cell types showed none or only 1-2 DEGs (Table S1), indicating that *BIN1* WT and KO neural cells have similar gene expression profiles. Importantly, even for the 41 DEGs observed in glutamatergic neurons, no significant enrichment for gene ontologies associated with endocytic pathway was observed (Table S2). These findings indicate that *BIN1* WT and KO iPSC-derived neurons mainly differ by the expression of BIN1. As a consequence, potential defects in endocytosis in these cell models are likely not due to major transcriptional modifications but due to intrinsic biological properties of BIN1 protein.

**Figure 6.**
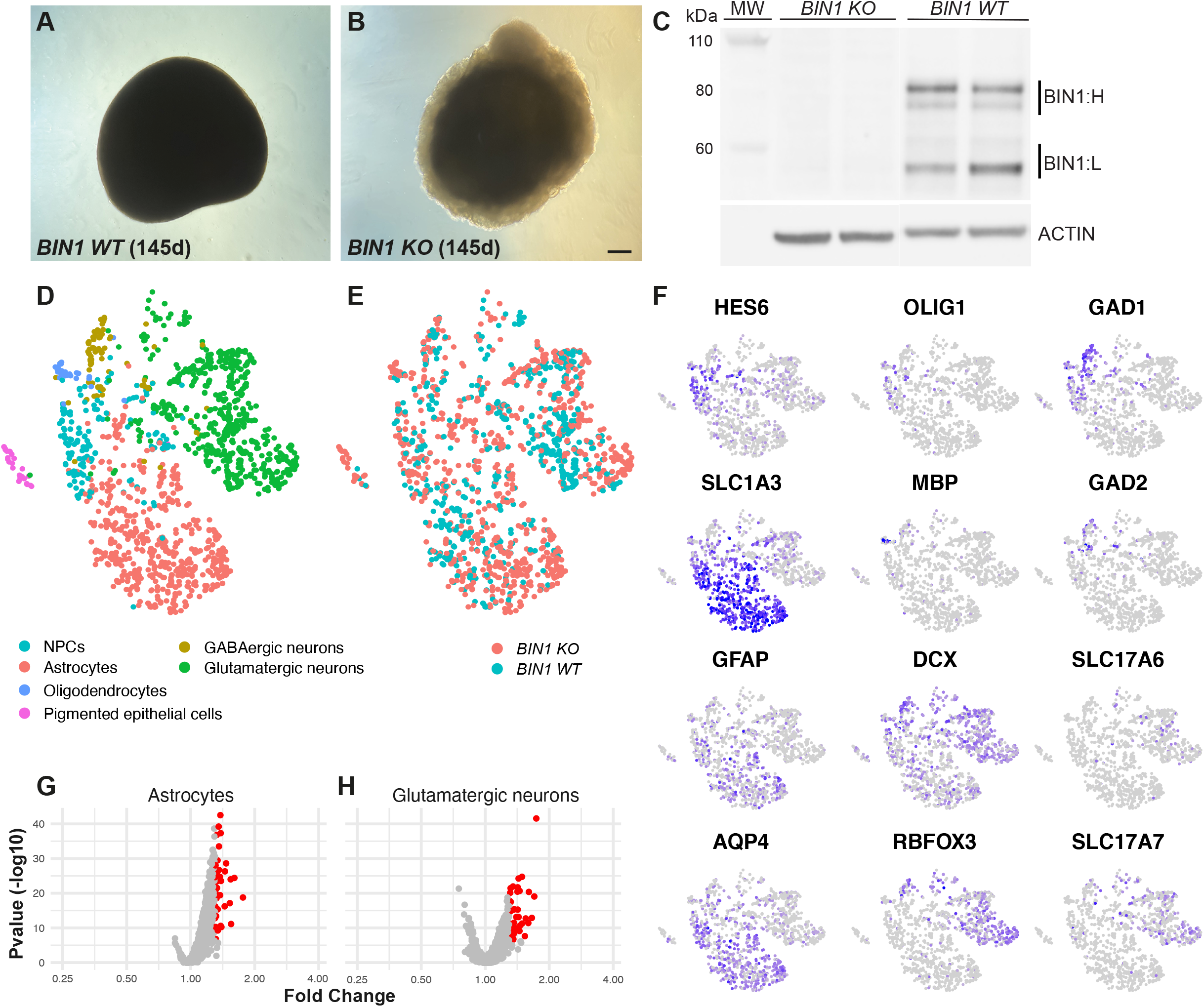
*BIN1* WT and KO cerebral organoids show similar composition and gene expression. **(A-B)** Brightfield images of a *BIN1* WT (A) and KO (B) cerebral organoids after 145 days (d) in culture. Calibration bar: 200µm. **(C)** Western blot for BIN1 and ACTIN in 190 days-old *BIN1* WT and KO cerebral organoids used for snRNA-seq. Bands corresponding to light (BIN1:L) and heavy BIN1 (BIN1:H) isoforms are indicated. MW – molecular weight. **(D-E)** tSNE plots of all 1441 cells from the snRNA-seq color-coded by cell type annotation (D) and genotype (E). **(F)** tSNE plots showing the expression of neural progenitor cell (HES6 and SLC1A3), astrocytes (SLC1A3, GFAP and AQP4), oligodendrocytes (OLIG1, MBP), pan neuronal (DCX, RBFOX3), GABAergic neurons (GAD1 and GAD2) and glutamatergic neurons (SLC17A6 and SLC17A7) markers. **(G-H)** Volcano plots showing genes differentially expressed in *BIN1* KO astrocytes (G) and glutamatergic neurons (H) compared to WT cells. Red dots indicate genes with fold change (FC) > 1.3 and false discovery rates (FDR) < 0.01.

### *BIN1* null mutation is associated with smaller early-endosome vesicles in hiPSC-derived neurons

In order to probe the impact of *BIN1* null mutation in iNs, we quantified the number and size of the early endosome antigen 1 (EEA1)-expressing endosomes in microtubule-associated protein 2 (MAP2)-expressing cells in both 2D cultures and cerebral organoids (Figure 7). EEA1 is an early endosomal Rab5 effector protein that has been implicated in the docking of incoming endocytic vesicles before fusion with early endosomes (Mu et al., 1995). We observed a significant change in the cumulative distribution of EEA1+ endosome volumes in hiNs both in 2D cultures and cerebral organoids, mostly due to a predominance of small volume endosomes in *BIN1* KO compared to WT iNs (Figure 7D and H). Conversely, no significant change in the number of endosomes was observed in iNs of both genotypes (Figure 7C and G). These data suggested that BIN1 is involved in the regulation of early endosome size in human neurons.

**Figure 7.**
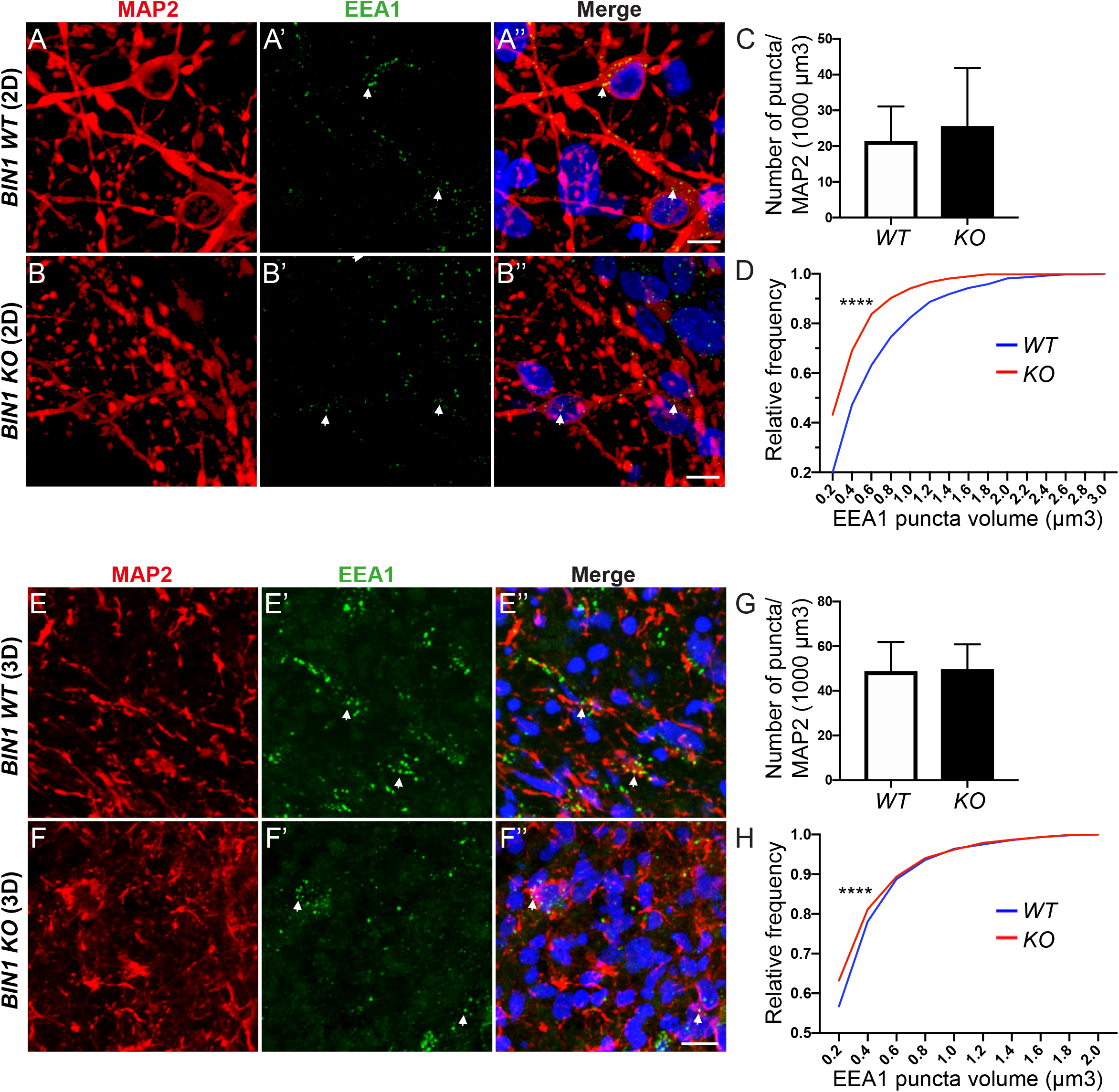
Increased proportion of small-volume endosomes in *BIN1* null mutant hiNs. **(A-B’’)** Representative images of *BIN1* WT and KO iNs in 6 weeks 2D cultures immunolabeled with antibodies against MAP2 (red, A and B) and EEA1 (green, A" and B") and stained with Hoechst 33258 (blue, A”and B”) (Scale Bar = 10 µm). **(C)** Plot showing the quantification of the number of EEA1+ puncta per 1000 µm^3^ of MAP2+ surface (N= 3 independent cell cultures; *BIN1 WT* iNs: 21.15 ± 9.94 EEA1+ puncta per 1000µm^3^ of MAP2+ surface; *BIN1 KO* iNs: 25.31 ± 16.58 EEA1+ puncta per 1000µm^3^ of MAP2+ surface; p=0.84, Unpaired t-test). **(D)** Plot showing the cumulative distribution of EEA1+ puncta volumes in *BIN1 WT* (blue line) and *KO* (red line) iNs (N= 3 independent cell cultures; Kolmogorov-Smirnov test, ****p<0.0001). **(E-F’’)** Coronal sections of 190 days old *BIN1 WT* and *KO* cerebral organoids immunolabeled with antibodies against MAP2 (red, E and F) and EEA1 (green, E" and F") and stained with Hoechst 33258 (blue, E”and F”). **(G)** Quantification of the number of EEA1+ puncta per 1000 µm^3^ of MAP2+ surface (N= 3 organoids per genotype; cerebral organoids: *BIN1 WT* iNs: 66.72 ± 18.13 EEA1+ puncta per 1000µm^3^ of MAP2+ surface; *BIN1 KO* iNs: 73.18 ± 12.47 EEA1+ puncta per 1000µm^3^ of MAP2+ surface; Unpaired t-test, p=0.96). **(H)** Cumulative distribution of EEA1+ puncta volumes in *BIN1 WT* (blue line) and *KO* (red line) iNs (N= 3 organoids per genotype; Kolmogorov-Smirnov test, ****p<0.0001).

### BIN1iso1 specifically modulates sizes of the early-endosome vesicles

We then wondered whether the function of BIN1 in endocytosis could also be isoform-specific in human neurons, as observed in Drosophila. To test this possibility, we set out to perform lentiviral-mediated transfection with fluorophore-expressing BIN1iso1, BIN1iso9 or control plasmids in iNs at 3 weeks of differentiation. After 3 additional weeks, we quantified the number and size of EEA1-expressing endosomes in transfected iNs identified by tdTomato expression (Figure 8). We observed that expression of BIN1iso1, but not BIN1iso9, in *BIN1* KO iNs fully rescued the volume of EEA1+ endosomes to values similar of those observed in WT iNs (Figure 8B). We also over-expressed BIN1iso1 and BIN1iso9 in WT iNs. Consistent with our previous observation in flies, we observed that BIN1iso1 overexpression in neurons led to an increase of large volumes EEA1+ endosomes, whereas the opposite effect was observed in neurons overexpressing BIN1iso9 (Figure 8C). Neither in *BIN1* WT nor in KO hiNs did we observe significant changes in the number of puncta per cell after BIN1iso1 or BINiso9 overexpression (Figure 8D). Lastly, we evaluated whether BIN1iso1 overexpression could have a neurotoxic effect in hiNs. To that end, we quantified the proportion of MAP2+ neurons in 6 weeks cultures after transduction with BIN1iso1-, BIN1iso9-or control-tdTomato+ cells. We found BIN1iso1 overexpression led to a 30% reduction in the proportion of neurons compared to controls (Figure 8E). Interestingly, this effect of BIN1iso1 was not observed in *BIN1* KO cells (data not shown), suggesting that only supra physiological expression levels of this isoform could be toxic for neurons.

**Figure 8.**
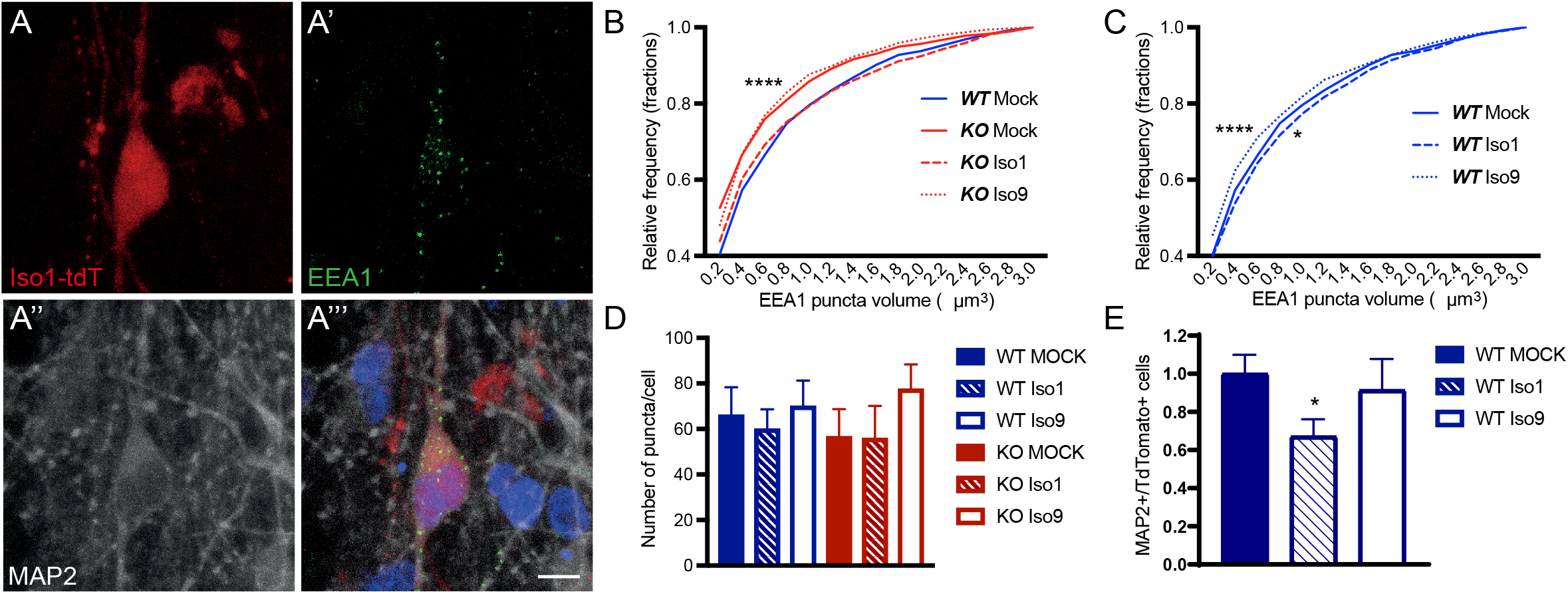
Rescue of endosomal phenotype in *BIN1* null mutant neurons by transduction of BIN1 Isoform 1. (A-A’’’) Representative images of *BIN1 WT* and *KO* iNs in 6-weeks-old 2D culture immunolabeled with antibodies against tdTomato (red, A), EEA1 (green, A’), MAP2 (grey, A”), and stained with Hoechst 33258 (blue, A’”) (Scale Bar = 10 µm). **(B)** Plot showing the frequency of occurrence of EEA1+ puncta of various sizes in transduced *BIN1 KO* iNs with tdT lentiviral constructs; *BIN1iso1* (dashed red line), *BIN1iso9* (dashed red line), and Mock-tdT (solid red line). *BIN1 WT* iNs tranduced with Mock-tdt (solid blue line) is indicated to show the rescue effect of the BIN1iso1 construct (orange line) in BIN1-null mutant neurons. Kolmogorov-Smirnov test followed by Bonferroni correction: *BIN1* WT+Mock vs *BIN1* KO+Mock: **** Padj<0.0001; *BIN1* WT +Mock vs *BIN1* KO + iso9: Padj<0.0001; *BIN1* KO + Mock vs *BIN1* KO + Iso1: **** Padj<0.0001; *BIN1* WT + Mock vs *BIN1* KO + Iso1: Padj=0.174; *BIN1* KO+ Mock vs *BIN1* KO + Iso9: Padj=0.207 (N=3 independent cell cultures). **(C)** Plot showing the frequency of occurrence of EEA1 puncta of various sizes in transduced *BIN1 WT* iNs with tdTomato (tdT)-expressing lentiviral constructs; *BIN1iso1* (dashed blue line), *BIN1iso9* (dotted blue line), and Mock-tdT (solid blue line). Kolmogorov-Smirnov test followed by Bonferroni correction: *BIN1* WT + Mock vs *BIN1* WT + Iso1: * Padj=0.04; *BIN1* WT + Mock vs *BIN1* WT + Iso9: **** Padj<0.0001 (N=3 independent cell cultures). **(D)** Graph showing the quantification of numbers of EEA1 puncta per cell in cells transduced with tdT-tagged lentiviral constructs (N=3 independent cell cultures. EEA1+ puncta/neuron: *BIN1 WT* + Mock: 65.86 ± 12.78; *BIN1 WT* + Iso1: 59.75 ± 8.895; *BIN1 WT* + Iso9: 69.82 ± 11.42; *BIN1 KO* + Mock: 56.47 ± 12.27; *BIN1 KO* + Iso1: 55.65 ± 14.52; *BIN1 KO* + Iso9: 77.31 ± 11.04; p=0.81, ANOVA _F(5,90)_). **(E)** Quantification of MAP2+/tdTomato+ neurons in BIN1iso1- and BIN1iso9-transduced cells relative to Mock-tdT-transduced cells (N=3 independent cell cultures; Number of tdT+ cells: Mock = 468; BIN1iso1 = 193; BIN1iso9 = 103; ANOVA followed by Dunnett"s multiple comparisons test, **p=0.0289).

Altogether, these observations indicate that BIN1iso1 expression is sufficient to regulate endosome volumes, even in the absence of other BIN1 isoforms in *BIN1* KO iNs, and that increased expression of BIN1iso1 deregulates endosome size and can lead to degeneration of human neurons, as observed in Drosophila photoreceptors.

## DISCUSSION

In this work, we assessed the function and potential neurotoxicity of human BIN1 isoforms in Drosophila and human neurons. We show a functional conservation of human BIN1 in Drosophila as BIN1iso1, BIN1iso8 and BIN1iso9 phenocopied dAmphA-induced photoreceptor neuron developmental defects and BIN1iso8 was able to rescue the locomotor defects associated with the loss of *dAmph* in Drosophila. Expression of BIN1iso1 resulted in a progressive neurodegeneration of photoreceptor neurons that was isoform-specific and dose-independent. The degeneration did not depend on the activation of the phototransduction cascade and was associated with a strong accumulation of vesicles harboring early and late endosomal markers. These results suggested an alteration of the endosome-lysosome pathway. Accordingly, the degeneration could be prevented by a loss of function of the early endosome regulator Rab5 and a gain of function of the recycling endosomal protein Rab11. Lastly, we observed a conservation of BIN1iso1 function in hiNs in 2D cultures and in organoids. Loss of BIN1 resulted in reduced size of early endosomes, whereas expression of BIN1iso1, but not BIN1iso9, was able to rescue the reduced size of early endosomes in the *BIN1* KO hiNs. As in Drosophila, overexpression of BIN1iso1 resulted in enlarged early endosomes and had neurotoxic effects in WT hiNs.

Our results indicate context and isoform-specific functions of BIN1: (i) Expression of BIN1iso1, BIN1iso8 and BIN1iso9 altered photoreceptor neuron rhabdomere morphogenesis during development in an isoform-independent and dose-dependent manner, whereas in the adult photoreceptor neurons BIN1iso1 expression induced neurodegeneration in an isoform-dependent and dose-independent manner; (ii) in human neurons, we observed a shift in BIN1 isoform expression from low molecular weight isoforms in iPSCs and NPCs (likely mainly BIN1iso9) to high molecular weight isoforms in hiNs (likely mainly BIN1iso1); (iii) BIN1iso1 expression rescued the size of BIN1-deficient early endosomes contrary to BIN1iso9. Collectively, these results argue for an important role of BIN1iso1 in mature neuronal cells and support the fact that BIN1iso1 is a neuron-specific isoform with specific functions in this cell-type (De Rossi et al., 2016; Zhou et al., 2014). Furthermore, (i) *BIN1* KO hiNs showed early endosomes of reduced size; as (ii) in *BIN1* KO hiNs, BIN1iso1 expression rescued early endosome sizes but did not lead to enlargement of these vesicles; and as (iii) in WT hiNs, BIN1iso1 overexpression induced early endosome enlargement. These results suggest that homeostatic BIN1iso1 expression levels need to be tightly regulated to allow a proper endocytic trafficking in neurons.

BIN1 has already been shown to regulate endocytosis (Calafate et al., 2016; Wigge et al., 1997) and endosome recycling (Ubelmann et al., 2017) as well as related processes in pre-or post-synaptic compartments (De Rossi et al., 2020; Schürmann et al., 2020). Our results suggest that BIN1iso1 functions in neurons take place at the early endosome crossroad as indicated (i) by the labeling of vesicles with early endosome markers in Drosophila, (ii) by the Rab5 rescue experiment in Drosophila and (iii) by the early endosome size modulation in hiNs. Additionally, in Drosophila, some of the vesicles also exhibited markers for late endosome/MVB and exceptionally lysosome. However, regulation of late endosome/MVB or lysosomal function did not modulate photoreceptor degeneration and secretion of exosomes did not appear to be blocked, suggesting that BIN1iso1 defect do not occur at this level. BIN1 has been proposed to regulate endocytosis by interaction with clathrin, Adaptor Proteins (AP) and dynamins (Dong et al., 2000; Grabs et al., 1997; Huser et al., 2013; Ramjaun and McPherson, 1998; Wigge et al., 1997). Since some of these proteins are also involved in vesicle budding of the intracellular organelles (Huser et al., 2013), BIN1iso1 may inhibit vesicle budding of early endosomes, thus, leading to an increase in their size. Of note, the phenotype with very large vesicles in drosophila likely results from Drosophila being a heterologous system for human BIN1iso1, and as a consequence, exhibiting stronger phenotypes.

Our results support a direct role of BIN1iso1 protein at the level of intracellular trafficking. Indeed, BIN1 knockout in cerebral organoids reduced endosome size without impacting expression of genes involved in the endosome trafficking pathway. In addition, our results in Drosophila showed that BIN1iso1 toxicity is dependent on the CLAP domain, which is known to bind directly intracellular trafficking proteins like clathrin and APs (Ramjaun and McPherson, 1998). Of note, BIN1iso9 overexpression in WT hiNs induced a reduction of early endosome size. Since BIN1 forms dimers through its BAR domain (Ramjaun et al., 1999), we postulate that BIN1iso9 may dimerize with BIN1iso1 and inhibit BIN1iso1 function in a dominant negative manner. Interestingly, this may also indicate another level of complexity to regulate BIN1 functions at the protein level.

*BIN1* is the second susceptibility gene for AD after *APOE* in terms of association level (Bellenguez et al., 2020) and interestingly, endosome enlargement has been described to be the first cytopathological marker of AD (Cataldo et al., 2000). It has also been shown that hiNs carrying mutations in APP or PSEN1, causal genes of familial AD, and hiNs depleted for the genetic risk factor SORL1 have enlarged early endosomes (Knupp et al., 2020; Kwart et al., 2019), similar to the phenotype observed in *BIN1 WT* hiNs after overexpression of BIN1iso1. This suggests that an increase in BIN1iso1 may be deleterious and contribute to early phases of the disease through early endosome enlargement. It implies an increase in BIN1iso1 levels in the disease, which remains highly debated with isoform-independent or isoform-related increased or decreased expression reported (Adams et al., 2016; Chapuis et al., 2013; Glennon et al., 2013; Holler et al., 2014; De Rossi et al., 2016; Taga et al., 2019, Marques-Coelho et al 2021). Moreover, most of the expression studies do not take into account disease progression and it is almost impossible to determine whether these expression variations may be a cause or a consequence of the disease development as previously mentioned. Finally, due to the many possible roles of BIN1 in AD pathophysiology, it is plausible that up- and/or down-regulation of expression of *BIN1* isoforms in the human brain would affect disease progression by different mechanisms, including endocytosis defects.

In conclusion, further complex experiments will be needed to better decipher how *BIN1* isoforms are involved in AD. Our results suggest that an increase of BIN1iso1 in neurons could contribute to AD pathogenesis by increasing the size of early endosomes observed early in the pathogenic process and by inducing neurodegeneration. Importantly, other AD genetic determinants have also been shown to regulate early endosome size, supporting early endosome defects as a major event in the pathogenesis of AD.

## Acknowledgements

The authors thank the BICeL platform of the Institut Biologie de Lille and the Vect"UB viral platform (INSERM US 005 – CNRS 3427 – TBMCore, Université de Bordeaux, France). The authors also thank the UMR 8199 LIGAN-PM Genomics platform (Lille, France) which belongs to the ‘Federation de Recherche’ 3508 Labex EGID (European Genomics Institute for Diabetes; ANR-10-LABX-46) and was supported by the ANR Equipex 2010 session (ANR-10-EQPX-07-01; ‘LIGAN-PM’). The LIGAN-PM Genomics platform (Lille, France) is also supported by the FEDER and the Region Nord-Pas-de-Calais-Picardie. Drosophila stocks obtained from the Bloomington Drosophila Stock Center (NIH P40OD018537) were used in this study. We thank A. Zelhof, G.L. Boulianne and J Laporte for antibody, fly stock and cDNA. The a5 and 4C5 antibodies developed by Fambrough, D.M. (The Johns Hopkins University) and de Couet, H.G. / Tanimura, T. (University of Hawaii) were obtained from the Developmental Studies Hybridoma Bank, created by the NICHD of the NIH and maintained at The University of Iowa, Department of Biology, Iowa City, IA 52242. Drosophila cDNA was obtained from the Drosophila Genomics Resource Center, supported by NIH grant 2P40OD010949. The Genotype-Tissue Expression (GTEx) Project was supported by the Common Fund of the Office of the Director of the National Institutes of Health, and by NCI, NHGRI, NHLBI, NIDA, NIMH, and NINDS. The data used for the analyses described in this manuscript were obtained from dbGaP accession number phs000424.v8.p2 on 04/02/2021. This work was supported by France Alzheimer (#328 ADhesion, #1999 BIN1DROSO), Fondation Vaincre Alzheimer (LECMA grant 13755) and la Fondation pour la recherche médicale (ALZ201912009628, ALZ201906008477). This work was co-funded by the European Union under the European Regional Development Fund (ERDF) and by the Hauts de France Regional Council (contrat no.18006176), the MEL (contract_2016_ESR_05), and the French State (contract no. 2018-3-CTRL_IPL_Phase2). This work was also funded by the Lille Métropole Communauté Urbaine and the French government"s LABEX DISTALZ program (Development of innovative strategies for a transdisciplinary approach to Alzheimer’s disease).

## Author contributions

Conceptualization, B.D., P.D., J.C.L., M.R.C. ; Methodology, P.D., M.R.C., X.H. ; Investigation, P.D., E.L., M.R.C., O.S., N.B., X.H., F.A., B.S.L., A.R.M.F., A.C., A.P., F.D. ; Resources, X.H., L.D., C.D., J.C., A.B. ; Writing - Original Draft, P.D., M.R.C., J.C.L., E.L., O.S. ; Writing - Reviews & Editing, P.D., M.R.C., J.C.L., E.L., O.S. ; Supervision, P.D., J.C.L., M.R.C., F.D., B.D., P.A., F.L., P.V. ; Funding Acquisition, J.C.L., M.R.C., P.D., B.D.

## Declaration of interests

The authors declare no competing interests.

## Methods

### Drosophila genetics and behavioral experiments

Flies were raised at 25°C under a light/dark cycle of 12h/12h (3000lux) on standard fly medium (Nutri-fly MF, Genesee Scientific, San Diego, CA, USA), unless otherwise stated. UAS-BIN1iso1, UAS-BIN1iso1 ΔEx7, UAS-BIN1iso1 ΔCLAP, UAS-BIN1iso9, UAS-BIN1iso8 and UAS-dAmphA lines were generated in this work (see supplementary methods). Briefly, cDNA were subcloned into pUASTattB vector and injected in attP2 lines (on the III chromosome) and in attP40 lines (on the II chromosome) (BestGene Inc., CA, USA). rh1-Gal4, GMR-Gal4, rh1-GFP, UAS-GFP:ninaC, UAS-evi:GFP, UAS-GFP:LC3 were described previously (Dourlen et al., 2012; Fouillet et al., 2012; Lauwers et al., 2018; Malmanche et al., 2017). The Amph^5E3^ line was a kind gift from GL Boulianne (Leventis et al., 2001). Other stocks were obtained from the Bloomington Drosophila Stock Center (BDSC, Bloomington, IN, USA): UAS-Luciferase (#35788), UAS-mCD8:GFP (#27400), UAS-GFP (#35786), Amph^MI08903-TG4.0^ (#77794), Rab5^EYFP^ (#62543), Rab7^EYFP^ (#62545), UAS-GFP-myc-2xFYVE (#42712), UAS-GFP-Rab5 (#43336), UAS-YFP.Rab5.Q88L (#9771), UAS-YFP.Rab5.S43N (#9774), Rab5^[2]^ (#42702), P{TRiP.HMS00147}attP2 Rab5 (#34832), UAS-Rab7.GFP (#42705), UASp-YFP.Rab7.Q67L (#24103), UASp-YFP.Rab7.T22N (#9778), UAS-Rab11-GFP (#8506), UASp-YFP.Rab11.Q70L (#9791), UASp-YFP.Rab11.S25N (#9792), P{TRiP.HMS01056}attP2 Vha68-2 (#64582), P{TRiP.HMS01442}attP2 VhaAC39-1 (#35029), UAS-mCherry:NLS (#38424), attP2 empty line (#8622), UASp-YFP.Rab9 (#9784), Rab1^EYFP^ (#62539), Rab6^EYFP^ (#62544), UAS-GFP.KDEL (#9898), UAS-ManII-EGFP (#65248), UAS-GFP-LAMP (#42714).

For the climbing test, 5 flies were subjected together to testing in a graduated cylinder. The wall of the cylinder had 5 main graduations, the top one being 13 cm from the bottom. Flies were tapped down to the bottom of the cylinder and recorded for 10 sec to see if they climbed up the wall of the cylinder. This was repeated 5 times in total. A score corresponding to the distance they had climbed was determined by the recorded movies. Flies got a score of between 0 to 5 depending on the main graduation that they were able to reach during the 10 sec period. The mean of the 5 trials was calculated and attributed to each fly.

### Western blot of Drosophila samples

Drosophila heads (n=10) and thorax (n=5) were dissected and crushed in ice-cold LDS lysis buffer (NP0008, NuPAGE, Novex, Life Technologies) supplemented with reducing agent (NP0009, NuPAGE, Novex, Life Technologies). Samples were centrifuged at 8500g for 10 min at 4°C. Supernatants were kept at -80°C. Once thawed, they were boiled for 10 min at 85°C before being loaded and separated in SDS–polyacrylamide gels 4–12% (NuPAGE Bis-Tris, ThermoScientific) in MOPS 1X buffer (NP0001-02, NuPAGE, Novex, Life Technologies). After migration, samples were transferred on to nitrocellulose membranes using the Biorad Trans-blot transfert system kit (Biorad) according to the supplier technical recommendation (7 min, 2.5 A, 25 V). Next, membranes were incubated in milk (5% in Tris-buffered saline with 0.1% Tween-20) to block non-specific binding sites during 1 h at RT, followed by several washes. Immunoblotting was carried out with primary antibodies anti-BIN1 (BIN1 99D, 05-449, Millipore, 1/2500, ab27796, abcam, 1/1000), anti-α-tubuline (α-tubuline DM1A, T9026, Sigma, 1/5000), anti-dAmph (#9906, kind gift of Andrew Zelhof, 1/5000)(Zelhof et al., 2001) and anti-GFP (anti-GFP, G1544, Sigma, 1/4000) overnight at 4°C. After washing, membranes were incubated with HRP-conjugated secondary antibodies (Jackson, anti-mouse 115-035-003 and anti-rabbit 111-035-003; ThermoScientific) 2 h at room temperature. Immunoreactivity was revealed using the ECL chemiluminescence system (WBLUC0500, Immobilon Classico Western HRP Substrate, Millipore) and imaged using the Amersham Imager 600 (GE LifeSciences). Optical densities of bands were quantified using Fiji software and results were normalized with respect to tubulin expression.

### Cornea neutralization

CO2-anesthetized flies were placed in a 35 mm cell culture dish half-filled with 1% agarose and covered with water at 4°C as described (Dourlen et al., 2013). Flies were observed using an upright confocal microscope (Zeiss LSM710, Wetzlar, Germany) equipped with a 40× water immersion long-distance objective. Images were acquired using the Zen acquisition software (Zeiss Zen software). Photoreceptor neurons were manually quantified.

### Immunofluorescence of Drosophila samples

Fly heads were dissected and fixed in 4% paraformaldehyde phosphate buffer saline (PBS) for 20 min at room temperature. After washing, retinas were finely dissected, permeabilized and depigmented in 0.3% (v/v) Triton X-100 in PBS (0.3% PBT) overnight at 4°C under gentle agitation. After blocking with 5% normal goat/donkey serum in 0.3% PBT, samples were incubated overnight at 4°C with the primary antibodies diluted in 0.3% PBT. The following antibodies were used: anti-NA/K ATPase alpha subunit (a5, DSHB, 1/100), anti-rhodopsin (4C5, DSHB, 1/200) and anti-GFP (132004, Synaptic System, 1/100). After washing they were incubated overnight at 4°C with Alexa 555 Phalloidin anti-F-actin (A34055, ThermoFisher Scientific) and the secondary antibodies diluted in 0.3% PBT: Alexa 488 Donkey anti-guinea pig (706-545-148, Jackson ImmunoResearch), Alexa 633 Goat anti-mouse (A-21052, ThermoFisher Scientific). After washing, samples were incubated in 90% glycerol PBS for 30 minutes in the dark before being mounted in the same solution. Retinas were imaged with a LSM710 confocal microscope (Zeiss, Wetzlar, Germany) equipped with a 40X oil objective.

### Electron microscopy

Drosophila eyes were dissected and fixed in 1% glutaraldehyde, 4% paraformaldehyde, 0.1M sodium cacodylate buffer (pH 6.8) 30 min at room temperature and then overnight at 4°C. After washing, eyes were post-fixed at room temperature in 1% OsO4 and 1.5% potassium ferricyanide for 1h, then with 1% uranyl acetate for 45 min, both in distilled water at room temperature in the dark. After washing, they were dehydrated with successive ethanol solutions. Eyes were infiltrated with epoxy resin (EMbed 812 from EMS) and were mounted in resin into silicone embedding molds. Polymerization was performed at 60°C for 2 days. Ultrathin sections of 70-80 nm thickness were observed on formvar-coated grid with a Hitachi H7500 TEM (Milexia, France), and images were acquired with a 1 Mpixel digital camera from AMT (Milexia, France).

### Maintenance of cells and generation of hiNPCs and mixed cultures of hiNs and hiAs

hiPSCs (ASE 9109, Applied StemCell Inc. CA, USA) modified for BIN1 in exon 3 (Figure 5) were generated by CRISPR/Cas9. Homozygous null mutants for BIN1 had a 5 bp deletion on one allele and an 8 bp deletion on the other allele. All hiPSCs, and all subsequent hiNPCs, hiNs, hiAs, and cerebral organoids derived thereof, were maintained in media from Stemcell Technologies, Vancouver, Canada. Maintenance of cell cultures and organoids were done in adherence with manufacturer"s protocols which can be found on the webpage of Stemcell Technologies. hiPSCs were maintained in mTeSR1 medium in non-treated cell culture dishes/plates pre-coated with vitronectin. Cell numbers and viability were recorded using a LUNA™ Automated Cell Counter.

In order to obtain hiNPCs, the embryoid body method detailed by Stemcell Technologies was used for the induction of BIN1 WT and KO hiPSCs. Following the generation of hiNPCs, these derived cells were maintained in treated cell culture dishes pre-coated with poly-L-ornithine (PLO) and laminin (5 µg/mL). PLO solution was made in water (0.001%) while laminin was diluted in PBS with Ca^2+^ and Mg^2+^. BIN1 WT and KO hiNPCs, thus generated, were maintained for up to 10 passages.

2D cultures comprising hiNs and hiAs were produced from hiNPCs. 60,000 hiNPCs/well were plated in 24-well cell imaging plates from Eppendorf (Cat # 0030741005) pre-coated with PLO (0.001%) and laminin (10 µg/mL). Cells were kept in 0.5 mL of NPC medium per well for 24 hours. Following this, equal volume of complete BrainPhys medium (supplemented with BDNF, GDNF, laminin, dibutyryl-cAMP, ascorbic acid, N2, and SM1) was added to each well to begin the process of differentiation. Subsequently, media was changed in the plates bi-weekly. The media change consisted of removing half of the existing medium in each well and replacing it with an equal volume of fresh complete BrainPhys medium. Mixed cultures of hiNs and hiAs were obtained at the end of 6 weeks from the start of the differentiation process.

### Generation of Cerebral Organoids

Cerebral Organoids were generated from hiPSCs at 80% confluency. hiPSCs were detached from the Vitronectin XF substrate using Gentle Cell Dissociation Reagent (Stem Cell Technologies), centrifuged, pelleted and resuspended in Embryoid Body (EB) seeding medium (Stem Cell Technologies) to form EBs. 9000 cells were plated per well in a 96-well round-bottom ultra-low attachment plate. After two days, 1 or 2 EBs were transferred to a well of a 24-well ultra-low attachment plate containing Induction Medium (Stem Cell Technologies). The EBs were kept in the induction Medium for 2 days and next they were transferred into Matrigel (Corning) using an embedding surface. When the Matrigel polymerized, the EB were transferred to a 6-well ultra-low adherent plate with Expansion Medium (Stem Cell Technologies). After 3 days, the medium was replaced by Maturation Medium (Stem Cell Technologies) and the plate was placed in an orbital shaker (65 rpm speed). Complete media changes were done on a bi-weekly basis.

### Lentiviral Infections

Lentiviral constructs were produced by the Vect"UB platform within the TBM Core unit at University of Bordeaux, Bordeaux, France (CNRS UMS 3427, INSERM US 005). All three lentiviral constructs harboured reporter tags for the expression of tdTomato protein. The lentiviral vectors used were the empty vector - 436 (ID # 1770), 436-Bin1Iso1 (ID # 1771), and 436-Bin1Iso9 (ID # 1772). Lentiviral infections were done in 3-week old differentiation cultures obtained from hiNPCs. Viral transductions were performed at a multiplicity of infection (MOI) of 1. In brief, appropriate volumes of each construct were mixed in complete BrainPhys medium and 50 ul of the viral medium mix was then added to each well. Each of the 3 constructs were added in triplicate to respective wells for each of the BIN1 WT and KO cells. Infected cells were maintained for a further 3-week period with bi-weekly changes of half volume of medium in each well.

### Immunocytochemistry of hiN samples

Cells were fixed in PFA (4% w/v) for 10 mins. Fixed cells were then washed with PBS 0.1 M. Cells were then blocked with blocking solution (5% normal donkey serum + 0.1% Triton X-100 in PBS 0.1 M) at room temperature for 1 hour under shaking conditions. Primary antibodies diluted in the blocking solution were then added and incubation was done overnight at 4°C under shaking conditions. The following day, cells were washed with PBS 0.1 M 3 times for 10 min. each before the addition of fluorophore-conjugated secondary antibodies in blocking solution for 2 hours at room temperature under shaking conditions ensuring protection from light. 3 washes with PBS were done for 10 min. each at room temperature under shaking conditions with protection from light. Hoechst 33258 nucleic acid stain was added to PBS 0.1 M in the second wash. Cells were mounted with fluoromount and imaged directly in the cell imaging plates. All images were acquired using an LSM 880 Confocal Scanning Microscope housed at the Imaging Platform of the Institut Pasteur de Lille using the ZEISS ZEN Imaging Software. Image acquisition was done at 40X for the various cellular markers in Figure 5. For the lentiviral transductions (Figure 7 and Figure 8), 10 images of different fields were acquired per well at 63X magnification using a zoom of 2.

### Immunohistochemistry of organoid samples

Cerebral organoids were fixed in PFA (4% w/v) for 30 min at 4°C followed by three washes with PBS 0.1 M. Cerebral organoids were then placed in sucrose solution (30% w/v) overnight before being embedded in O.C.T (Tissue-Tek). Embedded tissue was sectioned at 20 μm using a cryostat and mounted slides were stored at −80°C until immunostaining was performed. Mounted tissue was removed from storage and warmed by placing at room temperature for 30 min. Tissue were rehydrated and washed with room temperature PBS 0.1 M 3 times for 5 mins. Slides were then washed once with PBS with 0.2% Triton X-100 for 15 min. Tissue was blocked using 10% of donkey serum in PBS 0.1 M for 1 h at room temperature. After blocking, primary antibodies were added to 0.2 % triton X-100 and 10% of donkey serum in PBS 0.1 M at appropriate dilutions and incubated overnight at 4°C. The next day, slides were washed with PBS 0.1 M 3 times for 5 min each with gentle shaking. Subsequently, slides were incubated with fluorophore-conjugated secondary antibodies in 0.2 % Triton X-100 and 10% of donkey serum in PBS 0.1 M for 2 h at room temperature in the dark. After secondary antibody incubation, slides were washed 3 times with PBS for 5 min with gentle shanking. Nuclei were visualized by incubating the tissue for 5 min with Hoechst stain in PBS 0.1 M. Sections were mounted using aqueous mounting medium (Polysciences). Images were acquired using an LSM 810 Confocal Scanning Microscope in concert with the ZEISS ZEN imaging software housed at the Imaging Platform of the Pasteur Institute, Lille.

### Antibodies used for immunocytochemistry/immunohistochemistry of hiNs and organoid samples

hiPSCS were detected with antibodies for SOX2 and SSEA4 using the Molecular Probes™ Pluripotent Stem Cell 4-Marker Immunocytochemistry Kit (Fischer Scientific). Antibodies used for immunocytochemistry/immunohistochemistry were EEA1 (610457, BD Biosciences), MAP2 (188006, Synaptic Systems), RFP (600-401-379, Rockland Immunochemicals, Inc., SOX2 (14-9811-82, Invitrogen), NESTIN (MAB5326, Millipore), and GFAP (AB5804, Millipore). All fluorophore-tagged secondary antibodies were sourced from Jacskon ImmunoResearch Europe Ltd.

### Immunoblotting of 2D cultures and cerebral organoid

Samples from the 2D cultures or brain organoids were collected in RIPA buffer containing protease inhibitors (Complete mini, Roche Applied Science, Penzberg, Germany) and sonicated two times at 60% - 70% during 10 s before use for the western blotting analyses. Protein quantification was performed using the BCA protein assay (Thermo Scientific). In total, 10 μg of protein from extracts were separated in SDS–polyacrylamide gels 4–12% (NuPAGE Bis-Tris, Thermo Scientific) and transferred to nitrocellulose membranes (Bio-Rad). Next, membranes were incubated in milk (5% in Tris-buffered saline with 0.1% Tween-20 – TTBS, or SuperBlock – Thermo Scientific) to block non-specific binding sites during 1 h at RT, followed by several washes with TTBS. Immunoblottings were carried out with primary antibodies anti-BIN1 (ab182562, Abcam) and anti-β-ACTIN (A1978, Sigma-Aldrich) overnight at 4 °C on 20 RPM agitation. The membranes were washed three times in TTBS, followed by incubation with HRP-conjugated secondary antibodies anti-mouse (115-035-003) and anti-rabbit (111-035-003, Jackson ImmunoChemicals, Inc.) overnight at 4 °C on 20 rpm. The membranes were washed three times in TTBS, and the immunoreactivity was revealed using the ECL chemiluminescence system (SuperSignal, Thermo Scientific) and imaged using the Amersham Imager 600 (GE Life Sciences). Optical densities of bands were quantified using the Gel Analyzer plugin in Fiji – ImageJ.

### Image Analysis using Imaris

The surface function on Imaris was used for detection of EEA1 puncta. Threshold for the detection of spots is based on background subtraction. Following the application of automatic threshold, EEA1 puncta were detected. A manual filter cut-off was applied to detect all puncta volumes above 0.1 µm^3^ for the 2D cultures and above 0.02 µm^3^ for the cerebral organoids. In order to detect the MAP2 surfaces, absolute intensity was used as the parameter using automatic threshold. A filter of maximum intensity of 1 was applied to the MAP2 channel. For the detection of EEA1 puncta on MAP2 and EEA1 puncta on MAP2+/tdTomato+ cells, filters for the standard deviation of intensity were applied in the MAP2 and tdTomato channels respectively. Volume information for the EEA1 puncta were collated from each acquired image as CSV files. The volumes were sorted using Miscrosoft Excel. We used a cut-off maximum volume of 10 µm^3^ for each field. The sorted volume data were then analyzed statistically using the GraphPad Prism software.

### snRNA-seq library preparation

Nuclei isolation and Hash-tagging with oligonucleotides steps were realized on ice with pre-cold buffers and centrifugations at 4 °C. BIN1 WT and KO organoids of 6 months (one by condition) were cut in 2 parts, washed twice with 1 ml of Deionized Phosphate Buffer Saline 1X (DPBS, GIBCO™, Fisher Scientific 11590476) and centrifuged 5 min at 300 g. Organoids pellets were resuspended in 500μl lysis buffer (Tris-HCL 10 mM, NaCl 10 mM, MgCl2 3 mM, Tween-20 0,1%, Nonidet P40 Substitute 0.1%, Digitonin 0.01%, BSA 1%, Invitrogen™ RNAseout™ recombinant ribonuclease inhibitor 0.04 U/μL). Multiple mechanical resuspensions and wrecking steps in this buffer were perform for a total lysis time of 10 min, 500 μl of washing buffer was added (Tris-HCL 10 mM, NaCl 10 mM, MgCl2 3 mM, Tween-20 0.1%, BSA 1%, Invitrogen™ RNAseout™ recombinant ribonuclease inhibitor 0.04 U/μL) and the lysis suspension was centrifuged 8 min at 500 g (used for all following centrifugation steps). Nuclei pellets were washed tree times with one filtration step by MACS pre-separation filter 20 μm (Miltenyi Biotec). Nuclei pellets were resuspended in 100 μL of staining buffer (DPBS BSA 2%, Tween-20 0.01%), 10 μL of Fc blocking reagent HumanTruStainFc™ (422302, Biolegend) and incubated 5 min at 4°C. 1μl of antibody was added (Total-Seq™-A0451 anti-Vertebrate Nuclear Hashtag 1 MAb414 for the WT and Total-Seq™-A0453 anti-Vertebrate Nuclear Hashtag 3 MAb414 10µg for the KO, 97284 and 97286 respectively, Biolegend) and incubated 15 min at 4°C. Nuclei pellets were washed tree times in staining buffer with one filtration step by MACS pre-separation filter 20 μm (Miltenyi Biotec) to a final resuspension in 300 μl of staining buffer for Malassez cell counting with Trypan blue counterstaining (Trypan Blue solution, 11538886, Fisherscientific). Isolated nuclei were loaded on a Chromium 10x Genomics controller following the manufacturer protocol using the chromium single-cell v3 chemistry and single indexing and the adapted protocol by Biolegend for the HTO library preparation. The resulting libraries were pooled at equimolar proportions with a 9 for 1 ratio for Gene expression library and HTO library respectively. Finally, the pool was sequenced using 100 bp paired-end reads on the Illumina NovaSeq 6000 system following the manufacturer recommendations.

### snRNA-seq analysis and differential expression analysis

UMI Count Matrices for gene expression and for HTO libraries were generated using the CellRanger software (10x Genomics). After filtering for low quality cells according to the number of RNA, genes detected, and percentage of mitochondrial RNA, and normalizing the HTO matrix using centered log-ratio (CLR) transformation, 2,990 cells were assigned back to their sample of origin using HTODemux function of the SeuratV3 R Package (Satijalab), resulting to 1,794 and 1,196 cells for BIN1 KO and WT, respectively. Then, Seurat Workflow with SCTransform normalization was used to cluster the cells according to their transcriptome similarities. Each cluster was annotated using cell type specific markers. Finally, differential expression analysis between BIN1 KO and WT cells within each identified cell type was performed using DESeq2 package (Love et al., 2014).

## Data availability

The data that support the current findings are available upon request.

## Notes

### Competing Interest Statement

The authors have declared no competing interest.

## References

Adams, S.L., Tilton, K., Kozubek, J.A., Seshadri, S., and Delalle, I. (2016). Subcellular Changes in Bridging Integrator 1 Protein Expression in the Cerebral Cortex During the Progression of Alzheimer Disease Pathology. J. Neuropathol. Exp. Neurol. 75, 779–790.

Andrew, R.J., De Rossi, P., Nguyen, P., Kowalski, H.R., Recupero, A.J., Guerbette, T., Krause, S. V., Rice, R.C., Laury-Kleintop, L., Wagner, S.L., et al. (2019). Reduction of the expression of the late-onset Alzheimer’s disease (AD) risk-factor BIN1 does not affect amyloid pathology in an AD mouse model. J. Biol. Chem. 294, 4477–4487.

Bartscherer, K., Pelte, N., Ingelfinger, D., and Boutros, M. (2006). Secretion of Wnt Ligands Requires Evi, a Conserved Transmembrane Protein. Cell 125, 523–533.

Bellenguez, C., Küçükali, F., Jansen, I., Andrade, V., Morenau-Grau, S., Amin, N., Grenier-Boley, B., Boland, A., Kleineidam, L., Holmans, P., et al. (2020). Large meta-analysis of genome-wide association studies expands knowledge of the genetic etiology of Alzheimer’s disease and highlights potential translational opportunities. MedRxiv 17, 10.

Calafate, S., Flavin, W., Verstreken, P., and Moechars, D. (2016). Loss of Bin1 Promotes the Propagation of Tau Pathology. Cell Rep. 17, 931–940.

Cataldo, A.M., Peterhoff, C.M., Troncoso, J.C., Gomez-Isla, T., Hyman, B.T., and Nixon, R.A. (2000). Endocytic pathway abnormalities precede amyloid beta deposition in sporadic Alzheimer’s disease and Down syndrome: differential effects of APOE genotype and presenilin mutations. Am. J. Pathol. 157, 277–286.

Chapuis, J., Hansmannel, F., Gistelinck, M., Mounier, A., Van Cauwenberghe, C., Kolen, K. V, Geller, F., Sottejeau, Y., Harold, D., Dourlen, P., et al. (2013). Increased expression of BIN1 mediates Alzheimer genetic risk by modulating tau pathology. Mol. Psychiatry 18, 1225–1234.

Crotti, A., Sait, H.R., McAvoy, K.M., Estrada, K., Ergun, A., Szak, S., Marsh, G., Jandreski, L., Peterson, M., Reynolds, T.L., et al. (2019). BIN1 favors the spreading of Tau via extracellular vesicles. Sci. Rep. 9, 9477.

Dong, J., Misselwitz, R., Welfle, H., and Westermann, P. (2000). Expression and purification of dynamin II domains and initial studies on structure and function. Protein Expr. Purif. 20, 314–323.

Dourlen, P., Bertin, B., Chatelain, G., Robin, M., Napoletano, F., Roux, M.J., and Mollereau, B. (2012). Drosophila fatty acid transport protein regulates rhodopsin-1 metabolism and is required for photoreceptor neuron survival. PLoS Genet. 8, e1002833.

Dunst, S., Kazimiers, T., von Zadow, F., Jambor, H., Sagner, A., Brankatschk, B., Mahmoud, A., Spannl, S., Tomancak, P., Eaton, S., et al. (2015). Endogenously Tagged Rab Proteins: A Resource to Study Membrane Trafficking in Drosophila. Dev. Cell 33, 351–365.

Fouillet, A., Levet, C., Virgone, A., Robin, M., Dourlen, P., Rieusset, J., Belaidi, E., Ovize, M., Touret, M., Nataf, S., et al. (2012). ER stress inhibits neuronal death by promoting autophagy. Autophagy 8, 915–926.

Gatz, M., Reynolds, C.A., Fratiglioni, L., Johansson, B., Mortimer, J.A., Berg, S., Fiske, A., and Pedersen, N.L. (2006). Role of genes and environments for explaining Alzheimer disease. Arch. Gen. Psychiatry 63, 168–174.

Glennon, E.B.C., Whitehouse, I.J., Miners, J.S., Kehoe, P.G., Love, S., Kellett, K.A.B., and Hooper, N.M. (2013). BIN1 is decreased in sporadic but not familial Alzheimer’s disease or in aging. PLoS One 8, e78806.

Grabs, D., Slepnev, V.I., Songyang, Z., David, C., Lynch, M., Cantley, L.C., and De Camilli, P. (1997). The SH3 domain of amphiphysin binds the proline-rich domain of dynamin at a single site that defines a new SH3 binding consensus sequence. J. Biol. Chem. 272, 13419– 13425.

Holler, C.J., Davis, P.R., Beckett, T.L., Platt, T.L., Webb, R.L., Head, E., and Murphy, M.P. (2014). Bridging integrator 1 (BIN1) protein expression increases in the Alzheimer’s disease brain and correlates with neurofibrillary tangle pathology. J. Alzheimer’s Dis. 42, 1221–1227.

Huser, S., Suri, G., Crottet, P., and Spiess, M. (2013). Interaction of amphiphysins with AP-1 clathrin adaptors at the membrane. Biochem. J. 450, 73–83.

Knupp, A., Mishra, S., Martinez, R., Braggin, J.E., Szabo, M., Kinoshita, C., Hailey, D.W., Small, S.A., Jayadev, S., and Young, J.E. (2020). Depletion of the AD Risk Gene SORL1 Selectively Impairs Neuronal Endosomal Traffic Independent of Amyloidogenic APP Processing. Cell Rep. 31.

Kunkle, B.W., Grenier-Boley, B., Sims, R., Bis, J.C., Damotte, V., Naj, A.C., Boland, A., Vronskaya, M., van der Lee, S.J., Amlie-Wolf, A., et al. (2019). Genetic meta-analysis of diagnosed Alzheimer’s disease identifies new risk loci and implicates Aβ, tau, immunity and lipid processing. Nat. Genet. 51, 414–430.

Kwart, D., Gregg, A., Scheckel, C., Murphy, E., Paquet, D., Duffield, M., Fak, J., Olsen, O., Darnell, R., and Tessier-Lavigne, M. (2019). A Large Panel of Isogenic APP and PSEN1 Mutant Human iPSC Neurons Reveals Shared Endosomal Abnormalities Mediated by APP β-CTFs, Not Aβ. Neuron.

Lambert, J.C., Ibrahim-Verbaas, C.A., Harold, D., Naj, A.C., Sims, R., Bellenguez, C., DeStafano, A.L., Bis, J.C., Beecham, G.W., Grenier-Boley, B., et al. (2013). Meta-analysis of 74,046 individuals identifies 11 new susceptibility loci for Alzheimer’s disease. Nat. Genet. 45, 1452–1458.

Lauwers, E., Wang, Y.-C., Gallardo, R., Van der Kant, R., Michiels, E., Swerts, J., Baatsen, P., Zaiter, S.S., McAlpine, S.R., Gounko, N. V., et al. (2018). Hsp90 Mediates Membrane Deformation and Exosome Release. Mol. Cell 71, 689-702.e9.

Leventis, P.A., Chow, B.M., Stewart, B.A., Iyengar, B., Campos, A.R., and Boulianne, G.L. (2001). Drosophila Amphiphysin is a post-synaptic protein required for normal locomotion but not endocytosis. Traffic 2, 839–850.

Love, M.I., Huber, W., and Anders, S. (2014). Moderated estimation of fold change and dispersion for RNA-seq data with DESeq2. Genome Biol. 15.

Malki, I., Cantrelle, F.X., Sottejeau, Y., Lippens, G., Lambert, J.C., and Landrieu, I. (2017). Regulation of the interaction between the neuronal BIN1 isoform 1 and Tau proteins – role of the SH3 domain. FEBS J. 284, 3218–3229.

Malmanche, N., Dourlen, P., Gistelinck, M., Demiautte, F., Link, N., Dupont, C., Vanden Broeck, L., Werkmeister, E., Amouyel, P., Bongiovanni, A., et al. (2017). Developmental Expression of 4-Repeat-Tau Induces Neuronal Aneuploidy in Drosophila Tauopathy Models. Sci. Rep. 7, 40764.

Marques-Coelho, D., Iohan, L. da C.C., Melo de Farias, A.R., Flaig, A., Letournel, F., Martin-Négrier, M.L., Chapon, F., Faisant, M., Godfraind, C., Maurage, C.A., et al. (2021). Differential transcript usage unravels gene expression alterations in Alzheimer’s disease human brains. Npj Aging Mech. Dis. 7.

Mu, F.T., Callaghan, J.M., Steele-Mortimer, O., Stenmark, H., Parton, R.G., Campbell, P.L., McCluskey, J., Yeo, J.P., Tock, E.P.C., and Toh, B.H. (1995). EEA1, an early endosome-associated protein. EEA1 is a conserved α-helical peripheral membrane protein flanked by cysteine “fingers” and contains a calmodulin-binding IQ motif. J. Biol. Chem. 270, 13503– 13511.

Nicot, A.-S., Toussaint, A., Tosch, V., Kretz, C., Wallgren-Pettersson, C., Iwarsson, E., Kingston, H., Garnier, J.-M., Biancalana, V., Oldfors, A., et al. (2007). Mutations in amphiphysin 2 (BIN1) disrupt interaction with dynamin 2 and cause autosomal recessive centronuclear myopathy. Nat. Genet. 39, 1134–1139.

Prokic, I., Cowling, B.S., and Laporte, J. (2014). Amphiphysin 2 (BIN1) in physiology and diseases. J. Mol. Med. 92, 453–463.

Ramjaun, A.R., and McPherson, P.S. (1998). Multiple amphiphysin II splice variants display differential clathrin binding: identification of two distinct clathrin-binding sites. J. Neurochem. 70, 2369–2376.

Ramjaun, A.R., Philie, J., De Heuvel, E., and McPherson, P.S. (1999). The N terminus of amphiphysin II mediates dimerization and plasma membrane targeting. J. Biol. Chem. 274, 19785–19791.

De Rossi, P., Buggia-Prévot, V., Clayton, B.L.L., Vasquez, J.B., van Sanford, C., Andrew, R.J., Lesnick, R., Botté, A., Deyts, C., Salem, S., et al. (2016). Predominant expression of Alzheimer’s disease-associated BIN1 in mature oligodendrocytes and localization to white matter tracts. Mol. Neurodegener. 11, 59.

De Rossi, P., Nomura, T., Andrew, R.J., Masse, N.Y., Sampathkumar, V., Musial, T.F., Sudwarts, A., Recupero, A.J., Le Metayer, T., Hansen, M.T., et al. (2020). Neuronal BIN1 Regulates Presynaptic Neurotransmitter Release and Memory Consolidation. Cell Rep. 30, 3520-3535.e7.

Sartori, M., Mendes, T., Desai, S., Lasorsa, A., Herledan, A., Malmanche, N., Mäkinen, P., Marttinen, M., Malki, I., Chapuis, J., et al. (2019). BIN1 recovers tauopathy-induced long-term memory deficits in mice and interacts with Tau through Thr348 phosphorylation. Acta Neuropathol. 138, 631–652.

Schürmann, B., Bermingham, D.P., Kopeikina, K.J., Myczek, K., Yoon, S., Horan, K.E., Kelly, C.J., Martin-de-Saavedra, M.D., Forrest, M.P., Fawcett-Patel, J.M., et al. (2020). A novel role for the late-onset Alzheimer’s disease (LOAD)-associated protein Bin1 in regulating postsynaptic trafficking and glutamatergic signaling. Mol. Psychiatry 25, 2000–2016.

Sottejeau, Y., Bretteville, A., Cantrelle, F.-X., Malmanche, N., Demiaute, F., Mendes, T., Delay, C., Alves Dos Alves, H., Flaig, A., Davies, P., et al. (2015). Tau phosphorylation regulates the interaction between BIN1"s SH3 domain and Tau"s proline-rich domain. Acta Neuropathol. Commun. 3, 58.

Taga, M., Petyuk, V.A., White, C., Marsh, G., Ma, Y., Klein, H.-U., Connor, S.M., Khairallah, A., Olah, M., Schneider, J., et al. (2019). BIN1 protein isoforms are differentially expressed in astrocytes, neurons, and microglia: neuronal and astrocyte BIN1 implicated in Tau pathology. BioRxiv.

Taga, M., Petyuk, V.A., White, C., Marsh, G., Ma, Y., Klein, H.U., Connor, S.M., Kroshilina, A., Yung, C.J., Khairallah, A., et al. (2020). BIN1 protein isoforms are differentially expressed in astrocytes, neurons, and microglia: neuronal and astrocyte BIN1 are implicated in tau pathology. Mol. Neurodegener. 15.

Ubelmann, F., Burrinha, T., Salavessa, L., Gomes, R., Ferreira, C., Moreno, N., and Guimas Almeida, C. (2017). Bin1 and CD2AP polarise the endocytic generation of beta-amyloid. EMBO Rep. 18, 102–122.

Wang, T., and Montell, C. (2007). Phototransduction and retinal degeneration in Drosophila. Pflugers Arch. 454, 821–847.

Wigge, P., Köhler, K., Vallis, Y., Doyle, C.A., Owen, D., Hunt, S.P., and McMahon, H.T. (1997). Amphiphysin heterodimers: potential role in clathrin-mediated endocytosis. Mol. Biol. Cell 8, 2003–2015.

Zelhof, A.C., Bao, H., Hardy, R.W., Razzaq, A., Zhang, B., and Doe, C.Q. (2001). Drosophila Amphiphysin is implicated in protein localization and membrane morphogenesis but not in synaptic vesicle endocytosis. Development 128, 5005–5015.

Zhang, J., Schulze, K.L., Robin Hiesinger, P., Suyama, K., Wang, S., Fish, M., Acar, M., Hoskins, R.A., Bellen, H.J., and Scott, M.P. (2007). Thirty-one flavors of Drosophila Rab proteins. Genetics 176, 1307–1322.

Zhou, Y., Hayashi, I., Wong, J., Tugusheva, K., Renger, J.J., and Zerbinatti, C. (2014). Intracellular Clusterin Interacts with Brain Isoforms of the Bridging Integrator 1 and with the Microtubule-Associated Protein Tau in Alzheimer’s Disease. PLoS One 9, e103187.

