## Supplementary information and figures for "The Alzheimer susceptibility gene *BIN1* induces isoform-dependent neurotoxicity through early endosome defects"

#### Generation and validation of transgenic drosophila lines expressing human BIN1 isoforms

To assess the role of BIN1 isoforms, we generated 3 transgenic Drosophila lines expressing 3 representative BIN1 isoforms, brain BIN1-1, muscular BIN1-8 and ubiquituous BIN1-9. As a control, we also generated transgenic Drosophila lines expressing the longest dAmph isoform, dAmphA. We used the UAS/Gal4 system (Brand and Perrimon, 1993)(Figure S1A) and ΦC31-mediated targeted insertion of transgenes to express identical levels of BIN1 isoforms (Groth et al., 2004). We used the attP40 and attP2 landing sites on the second and third chromosomes. We got 1 to 5 lines for each construct resulting from independent identical insertion events. To test the expression of human BIN1 isoforms, we crossed the lines with an eye-specific GMR driver line and performed a western blot analysis. The BIN1 isoforms were expressed with an expected difference in the molecular weight (BIN1-1 theoretical MW 65kDa > BIN1-8 theoretical MW 50kDa > BIN1-9 theoretical MW 48kDa). Surprisingly, some of the supposed-to-be identical lines expressed different levels of the same isoform. We selected 2 lines of each isoform inserted on the third chromosome and tested their expression at the RNA level by RT-qPCR (Figure S1B). We observed that basal expression levels were the same for all isoforms (BIN1-1#3, BIN1-8#1, BIN1-8#2, BIN1-9#2) except for two lines (BIN1-1#1, BIN1-9#1) expressing around twice as much BIN1 isoforms. This is likely due to the insertion of 2 copies of the transgenes. We repeated the western blot analysis for these lines which confirmed the higher expression of the two lines at the protein levels (Figure S1C). We decided to keep and use these lines to test for dose-dependent effects. We also observed that the basal level-expressing BIN1-1 line exhibited non-significant higher protein levels of BIN1-1 than BIN1-8 and BIN1-9 (significant difference only between BIN1-1#3 and BIN1-8#1, $p=0.023$, ANOVA with post-hoc Tukey test). This indicates that BIN1-1 tended to be more stable than BIN1-8 and BIN1-9 when expressed in the Drosophila eye. In the article, if not specified, we have used the basal level-expressing lines (BIN1-1#3, BIN1-8#1, BIN1-9#2) on the third chromosome and “BIN1-1/BIN1-9 high” refers to the BIN1-1#1 and BIN1-9#1 highly expressing lines.

#### Characterization of dAmphMI08903-TG4.0 allele

The dAmph$^{MI08903\text{-}TG4.0}$ allele contains a Trojan Gal4 exon cassette in the first intron of dAmph (Diao et al., 2015; Lee et al., 2018) (Figure S2). This allows Gal4 expression under the control of dAmph endogenous promoter while arresting dAmph transcription thanks to a polyadenylation signal located 3’ of the GAL4. Because the first exon only contains 68nt of the dAmph coding DNA sequence, the expressed truncated protein consists only in around the first 25 amino acids of the protein and the mutation is likely null. To test this, we assessed dAmph protein expression and the climbing ability of compound heterozygous dAmph$^{MI08903\text{-}TG4.0}$ flies with the known dAmph$^{5E3}$ null allele (Leventis et al., 2001). We did not detect any dAmph in dAmph$^{MI08903\text{-}TG4.0/5E3}$ compound heterozygous fly protein extract (Fig. 1b). In addition, these flies had strong locomotor defect. They had a climbing score close to the one of Amph$^{5E3/5E3}$, around 2, whereas control flies had a climbing score close to 5 (Figure 1C). These results indicate that the Amph$^{MI08903\text{-}TG4.0}$ allele can be considered as a null allele.

### Supplementary methods

#### Construction of Drosophila transgenesis vectors

BIN1iso1 cDNA, BIN1iso8 cDNA (kind gifts of J Laporte) and BIN1iso9 cDNAs (SC128163, OriGene Technologies, Inc., USA) were amplified by PCR using the forward primer CAAAATGGCAGAGATGGGCAGTAA and the reverse primer CTCGAGTCATGGGACCCTCTCAGTG, which allowed the insertion of a CAAA Kozak sequence upstream of the ATG and a CTCGAG XhoI restriction enzyme site after the stop codon. Of note, BIN1iso9 cDNA has two synonymous mutations, a C instead of a T at position 486 and a T instead of a C at position 864 from the initial ATG. Similarly dAmphA cDNA (LD19810 from the Drosophila Genomics Resource Center, Indiana University) was amplified by PCR using the forward primer CAAAATGACCGAAAATAAAGGCATAA and the reverse primer ACTTCACGCGTCCCATCTGACTCGAG. The PCR product was cloned into a pGEM-T Easy Vector (Promega). After sequence checking, we subcloned the insert using EcoRI and XhoI restriction enzyme into a pUAST-attB transgenesis vector. After sequence checking, transgenesis plasmids were sent to the Bestgene company (BestGene Inc, Chino Hills, USA) for embryo injection in y1 w67c23; P{CaryP}attP40 and $y^{1}$ $w^{67c23}$;; P{CaryP}attP2 lines (attP sites on the II and III chromosomes respectively).

For BIN1iso1 ΔEx7 and BIN1iso1 ΔCLAP, cDNAs were synthetized by Invitrogen GeneArt gene synthesis adding upstream an EcoRI restriction enzyme site from the pGEMT-easy and the CAAA Kozak sequence, and adding downstream an XhoI site to have identical sequences around the cDNA and to allow the comparison with BIN1iso1 transgenic flies. BIN1iso1 ΔEx7 and BIN1iso1 ΔCLAP cDNAs correspond to BIN1iso1 cDNA without Exon7 for the former and Exon13-14-15-16 for the latter. cDNA was subcloned into a pUAST attB transgenesis vector and transgenic flies were generated as described above.

#### RT-qPCR

RNA was extracted from 30 Drosophila heads per condition using Trizol reagent (15596018, Invitrogen) according to manufacturer instructions. Retrotranscription was performed using ThermoScript™ RT-PCR System (11146-024, Invitrogen) and PCR was performed using … in the AriaMx Real-time PCR System (G8830A, Agilent Technologies). The following primers were used: for BIN1, forward primer ATGTTCAAGGTACAGGCCCA and reverse primer TCAGTGAAGTTCTCGGGGAA, both primers are localized in exons common to all isoforms corresponding to the SH3 domain; for RpL32, forward primer CCAAGGACTTCATCCGCCACC and reverse primer GCGGGTGCGCTTGTTCGATCC.

### References

Brand, A.H., and Perrimon, N. (1993). Targeted gene expression as a means of altering cell fates and generating dominant phenotypes. Development *118*, 401–415.

Diao, F., Ironfield, H., Luan, H., Diao, F., Shropshire, W.C., Ewer, J., Marr, E., Potter, C.J., Landgraf, M., and White, B.H. (2015). Plug-and-play genetic access to drosophila cell types using exchangeable exon cassettes. Cell Rep. *10*, 1410–1421.

Groth, A.C., Fish, M., Nusse, R., and Calos, M.P. (2004). Construction of Transgenic Drosophila by Using the Site-Specific Integrase from Phage ϕC31. Genetics *166*, 1775–1782.

Lee, P.T., Zirin, J., Kanca, O., Lin, W.W., Schulze, K.L., Li-Kroeger, D., Tao, R., Devereaux, C., Hu, Y., Chung, V., et al. (2018). A gene-specific T2A-GAL4 library for drosophila. Elife *7*.

Leventis, P.A., Chow, B.M., Stewart, B.A., Iyengar, B., Campos, A.R., and Boulianne, G.L. (2001). Drosophila Amphiphysin is a post-synaptic protein required for normal locomotion but not endocytosis. Traffic *2*, 839–850.

**Figure S1**: **Creation of transgenic lines expressing human BIN1 isoforms**. **(A)** Scheme of the Gal4/UAS system with the example of a GMR driver and a UAS construct expressing BIN1. **(B)** RT-qPCR analysis of BIN1 isoforms for 2 lines of each isoforms on the third chromosome (ANOVA, post-hoc Tukey, * p<0.05, **** p<0.0001). **(C)** Western blot analysis of BIN1 isoforms for the same lines. Tubulin is used as a loading control (ANOVA, post-hoc Tukey, * p<0.05, ** p<0.01, **** p<0.0001).

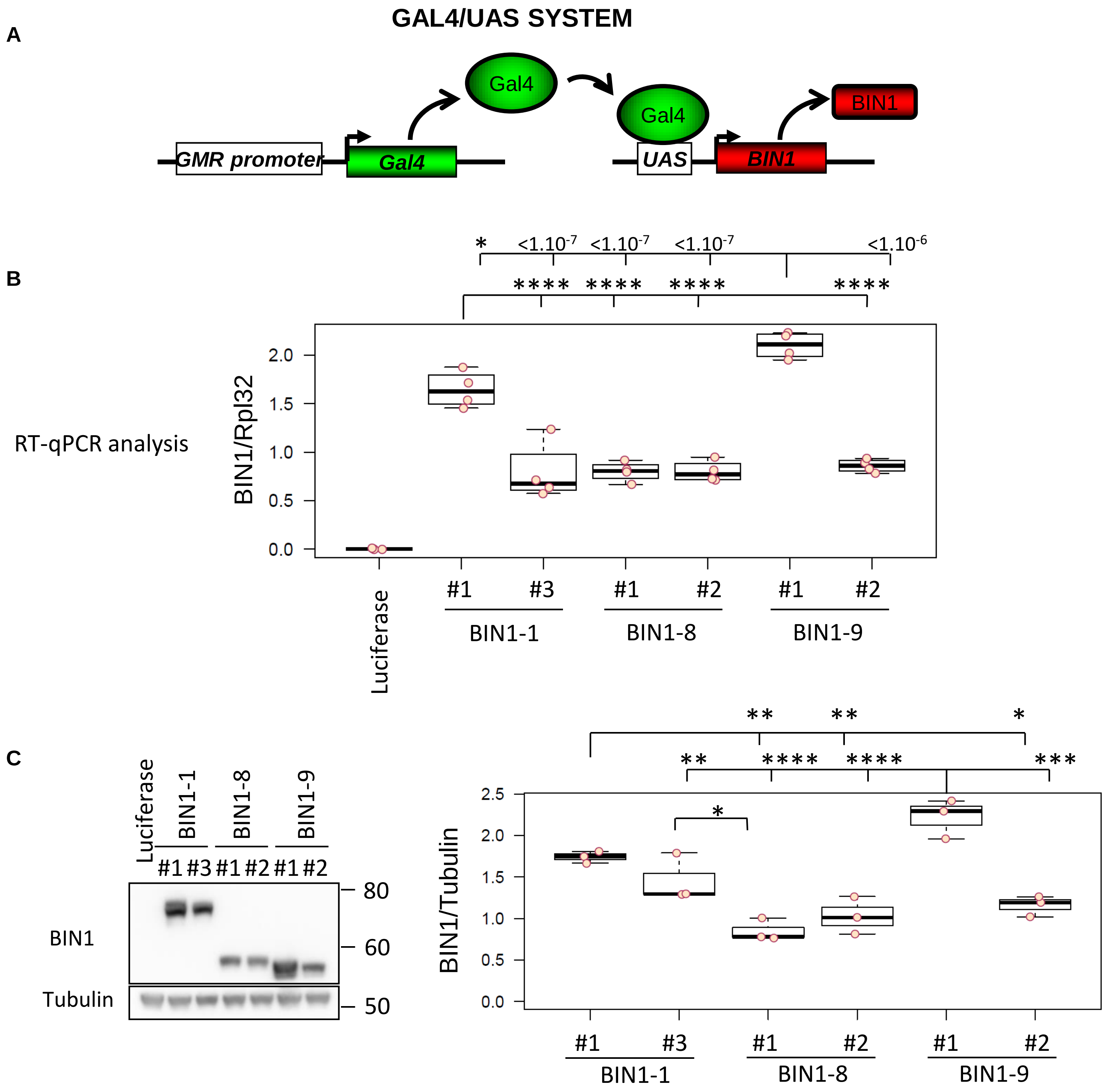

**Figure S2**: Scheme of Amph$^{MI08903\text{-}TG4.0}$ allele. SA=Splice acceptor site, SD=Splice donnor site.

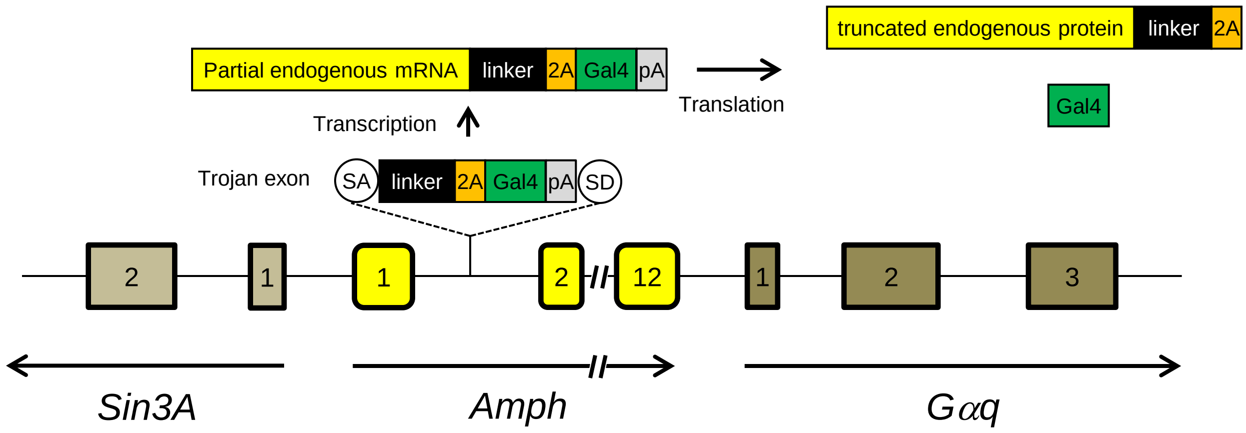

**Figure S3 : Generation and test of transgenic Drosophila expressing truncated human BIN1-1 forms for the Exon7 (BIN1-1 ΔEx7) and the CLAP domain (BIN1-1 ΔCLAP).** The two UAS constructs were inserted in the attP40 (2nd chromosome) and the attP2 (3rd chromosome) landing sites thanks to the ΦC31 integrase. We obtained 3 to 5 independent lines per landing sites. **(A)** Western blot analysis of the transgene expression under rh1 driver and quantification **(B)**. Each line expressed a BIN1 form with the expected molecular weight (BIN1-1 > BIN1-1 ΔEx7 > BIN1-1 ΔCLAP > BIN1-9). Expression levels were similar between lines with a non-significant tendency of decreased levels for BIN1-1 ΔCLAP. Two lines of each category were tested for their ability to induce photoreceptor neuron degeneration. **(C)** Quantification of the truncated BIN1-1 form-induced neurodegeneration. Results were identical whatever the insertion sites and independent lines. Loss of Exon7 partially rescued photoreceptor neurons whereas loss of the CLAP nearly totally rescued them. We can wonder if the reduced expression of the BIN1-1 ΔCLAP contribute to its lesser toxicity but we have seen with BIN1-1 and BIN1-9 that the effect is likely dose-independent. Results of BIN1-1 ΔEx7 attP2 #1 and of BIN1-1 ΔCLAP attP2 #1 were used in Figure 2F and 2G.

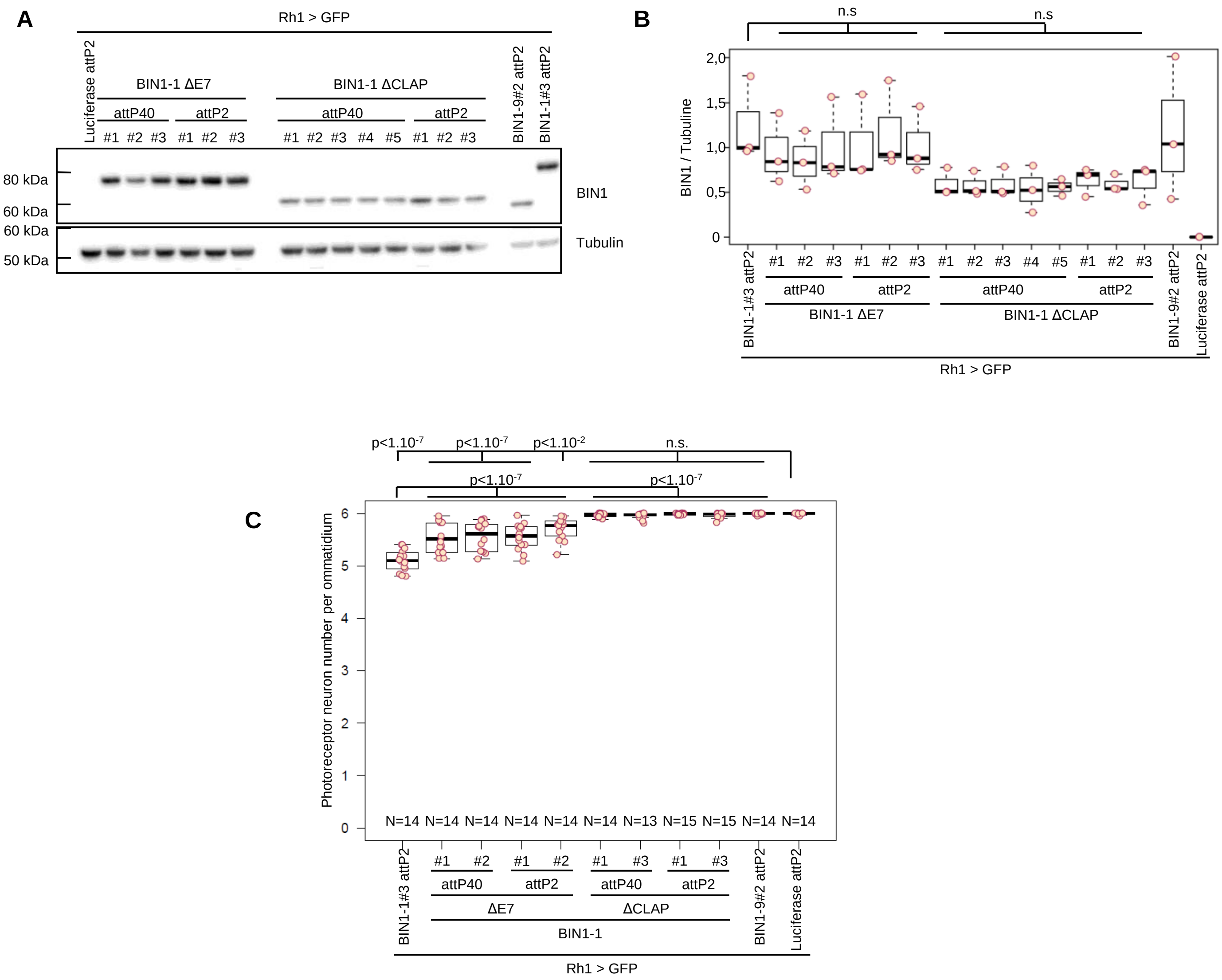

**Figure S4 : Electron microscopy images of BIN1-1-induced photoreceptor degeneration in Drosophila eyes. (A)** View of several ommatidia expressing luciferase as a control (left) or BIN1-1 (right) in 15 day-old flies. BIN1-1 expressing ommatidia exhibit many vesicles. **(B)** Longitudinal view of photoreceptor neurons in the distal part of the retina. Under the cornea (¤) and pseudocone (*), a photoreceptor neuron is filled with vesicles (arrow). The nucleus (arrow in the first inset) is squeezed but the chromatine seem normal. The cytoplasm is pushed on the side against the plasma membrane, which is not disrupted (arrow in the second inset). **(C)** Image showing a multilamellar body.

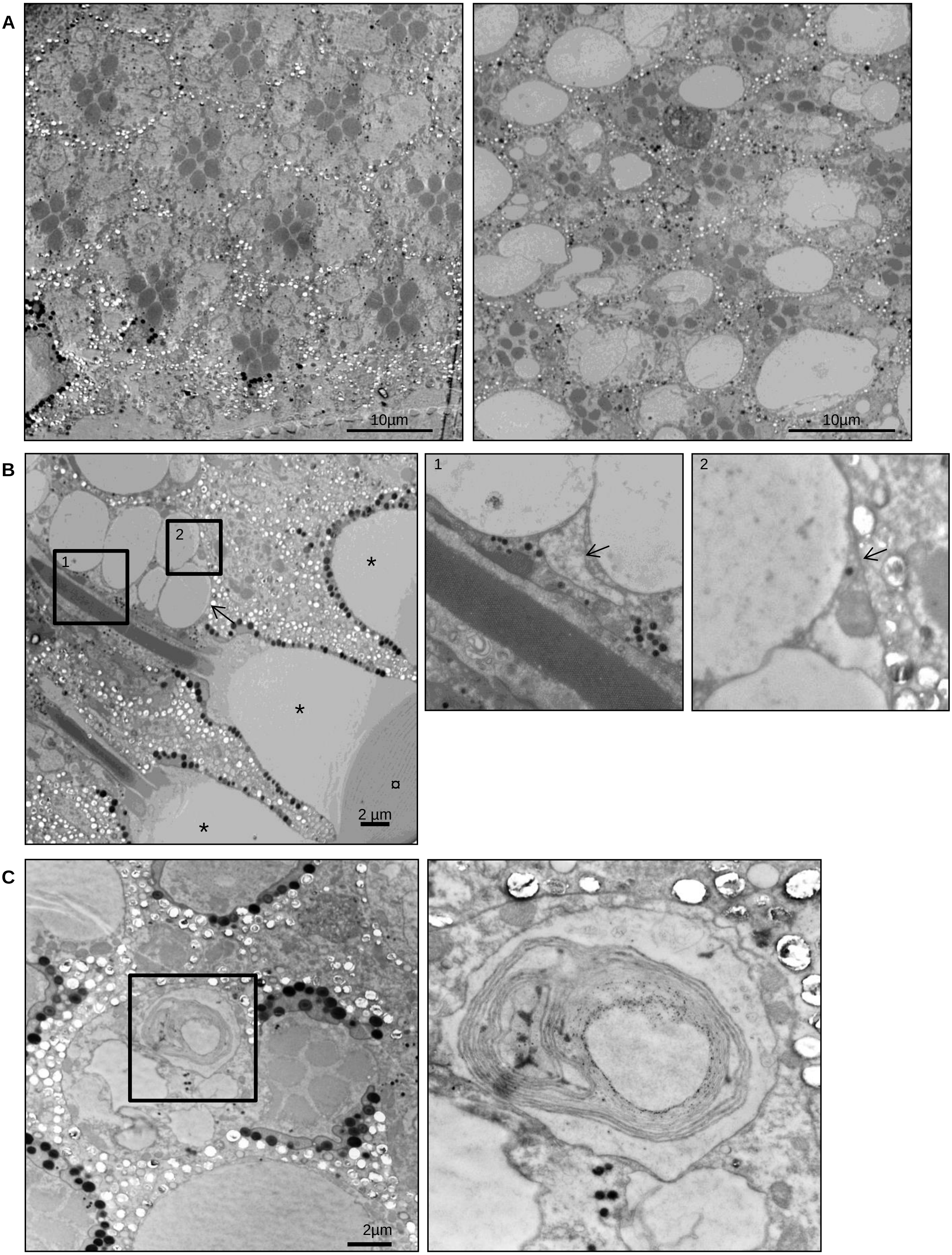

**Figure S4 : Electron microscopy images of BIN1-1-induced photoreceptor degeneration in Drosophila eyes. D** BIN1-1 expressing retina exhibited dying photoreceptor neurons. They first started to round up, their cytoplasm became electron-dense with abnormal mitochondria (arrow, upper left panel). They shrank (arrow, upper right panel) and were finally phagocytosed by the adjacent interommatidial cell (arrow, lower left panel).

D

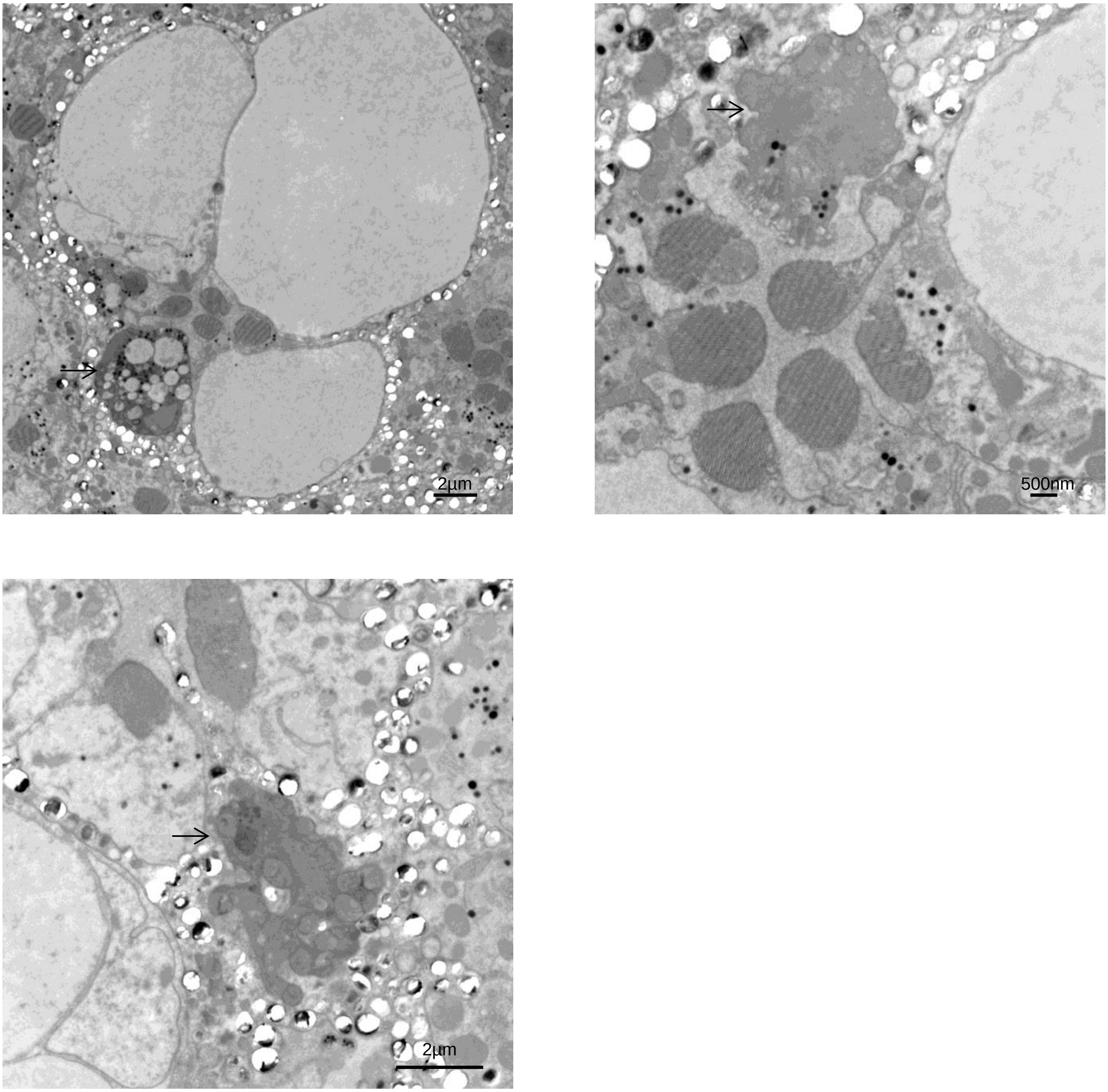

**Figure S5: Screening of organelle markers in BIN1-1-induced degenerating photoreceptor neurons.** BIN1-1-expressing flies were crossed with line expressing green fluorescent marker for ER **(A)**, Golgi **(B)**, plasma membrane **(C)**, early endosome **(D)**, late endosome/multivesicular body **(E)**, recycling endosome **(F)**, lysosome **(G)** and autophagosome **(H)**, and we let flies age for 1 and 2 weeks before dissection and immunofluorescence. Rh1 and actin labelling (respectively white and red in merge images) were used to visualize retinal structure and only green channels and merge images are shown. Note that the KDEL:GFP marker labelled nuclear envelop (arrows in **A**), early and late endosome/multivesicular body marker labelled small to middle size vesicles (arrows in **D**, **E**), the evi:GFP marker also labelled bigger vesicles (see image of one week-old flies) and the interrhabdomeric space (arrowhead in **E**), which likely corresponds to released exosomes, the Lamp2:GFP marker labelled on rare cases some middle to big size vesicles (arrow in **G**) and the autophagosome marker GFP:LC3 labelled small structures in the control and BIN1-1 conditions (arrows in **H**).

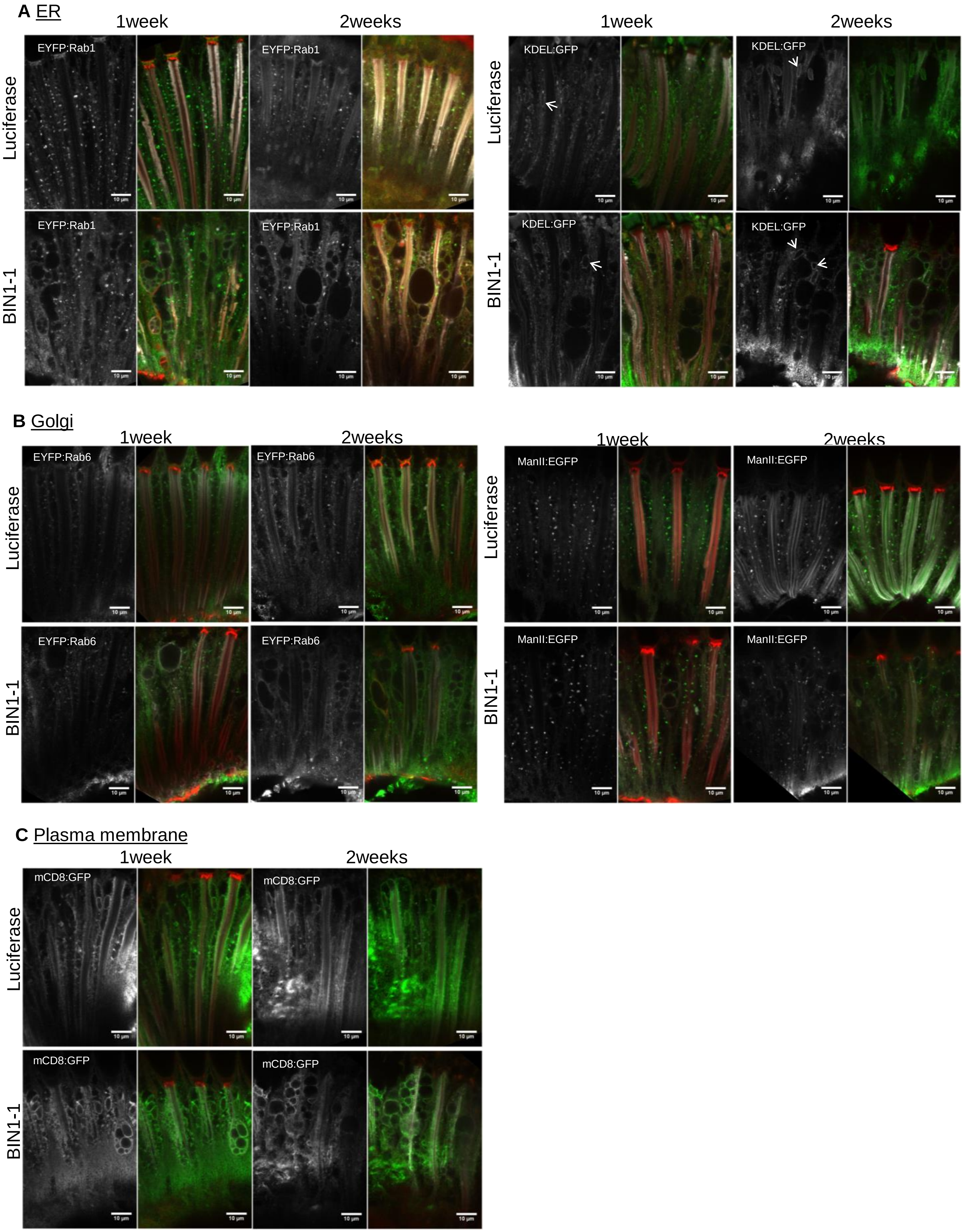

**Figure S5: Screening of organelle markers in BIN1-1-induced degenerating photoreceptor neurons.** BIN1-1-expressing flies were crossed with line expressing green fluorescent marker for ER **(A)**, Golgi **(B)**, plasma membrane **(C)**, early endosome **(D)**, late endosome/multivesicular body **(E)**, recycling endosome **(F)**, lysosome **(G)** and autophagosome **(H)**, and we let flies age for 1 and 2 weeks before dissection and immunofluorescence. Rh1 and actin labelling (respectively white and red in merge images) were used to visualize retinal structure and only green channels and merge images are shown. Note that the KDEL:GFP marker labelled nuclear envelop (arrows in **A**), early and late endosome/multivesicular body marker labelled small to middle size vesicles (arrows in **D**, E), the evi:GFP marker also labelled bigger vesicles (see image of one week-old flies) and the interrhabdomeric space (arrowhead in **E**), which likely corresponds to released exosomes, the Lamp2:GFP marker labelled on rare cases some middle to big size vesicles (arrow in **G**) and the autophagosome marker GFP:LC3 labelled small structures in the control and BIN1-1 conditions (arrows in **H**).

**D** Early endosome

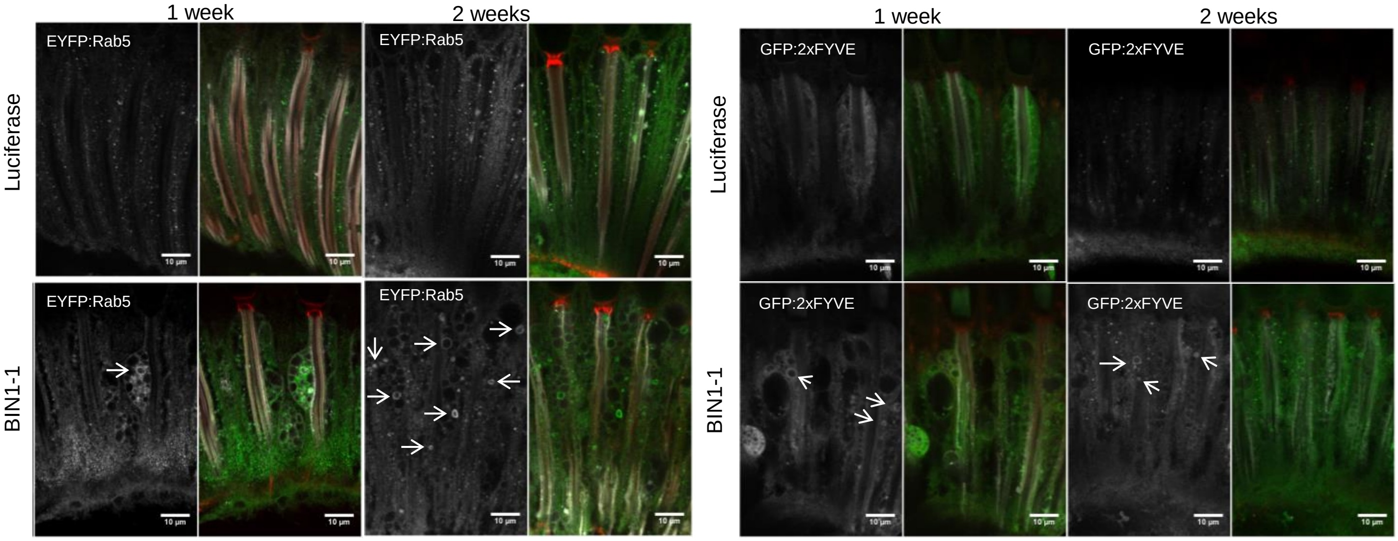


**E** Late endosome

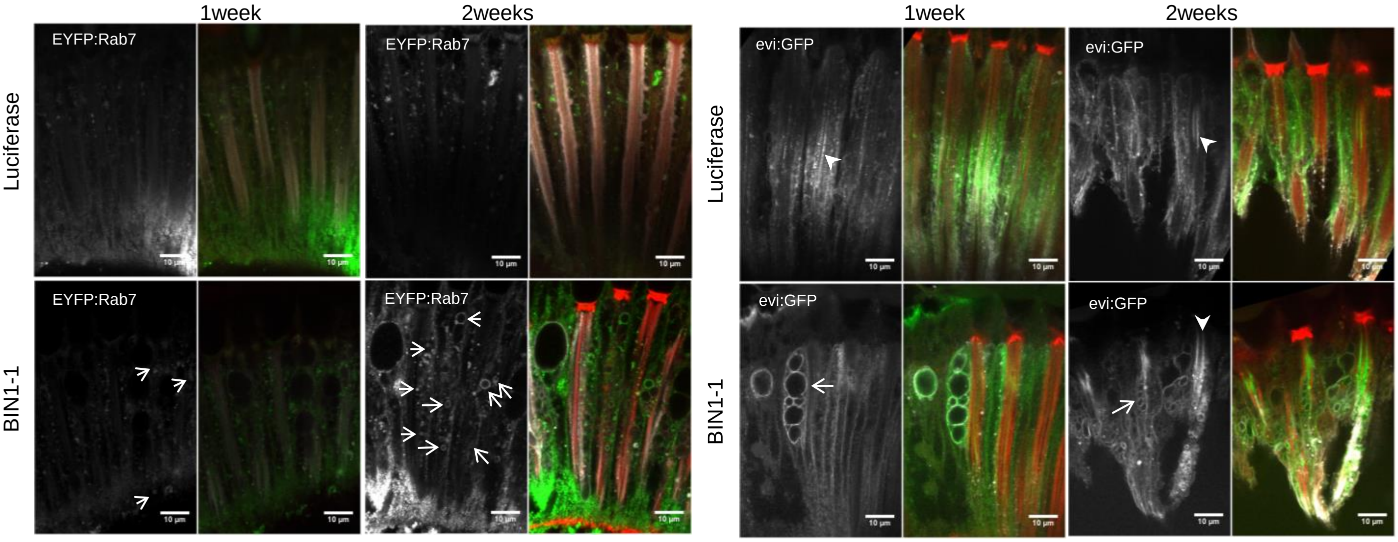


**F** Recycling endosome

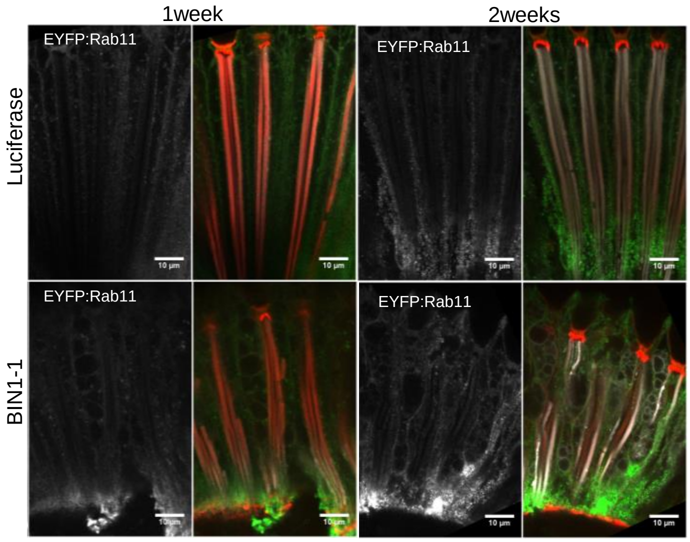

**Figure S5: Screening of organelle markers in BIN1-1-induced degenerating photoreceptor neurons.** BIN1-1-expressing flies were crossed with line expressing green fluorescent marker for ER **(A)**, Golgi **(B)**, plasma membrane **(C)**, early endosome **(D)**, late endosome/multivesicular body **(E)**, recycling endosome **(F)**, lysosome **(G)** and autophagosome **(H)**, and we let flies age for 1 and 2 weeks before dissection and immunofluorescence. Rh1 and actin labelling (respectively white and red in merge images) were used to visualize retinal structure and only green channels and merge images are shown. Note that the KDEL:GFP marker labelled nuclear envelop (arrows in **A**), early and late endosome/multivesicular body marker labelled small to middle size vesicles (arrows in **D**, E), the evi:GFP marker also labelled bigger vesicles (see image of one week-old flies) and the interrhabdomeric space (arrowhead in **E**), which likely corresponds to released exosomes, the Lamp2:GFP marker labelled on rare cases some middle to big size vesicles (arrow in **G**) and the autophagosome marker GFP:LC3 labelled small structures in the control and BIN1-1 conditions (arrows in **H**).

**G** Lysosome

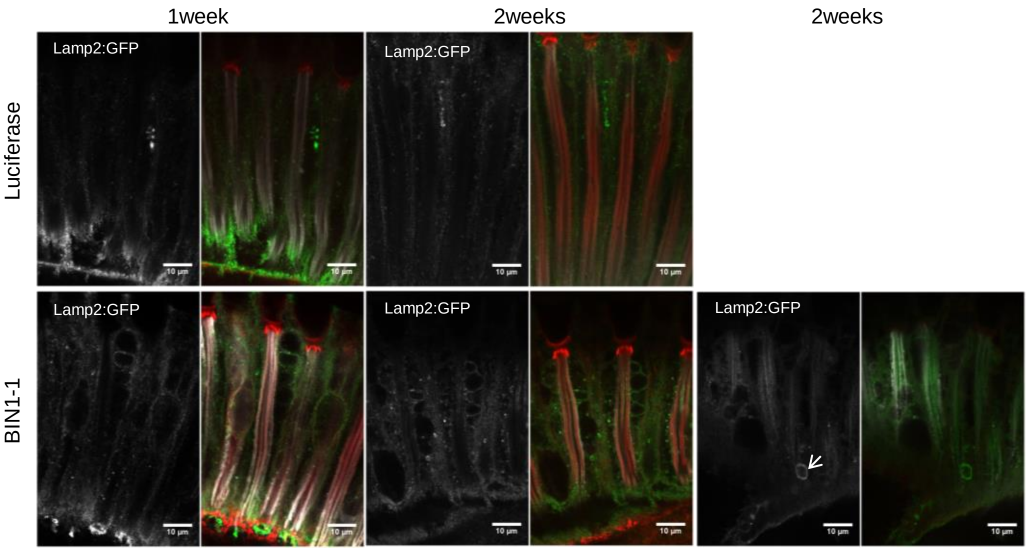


**H** autophagosome

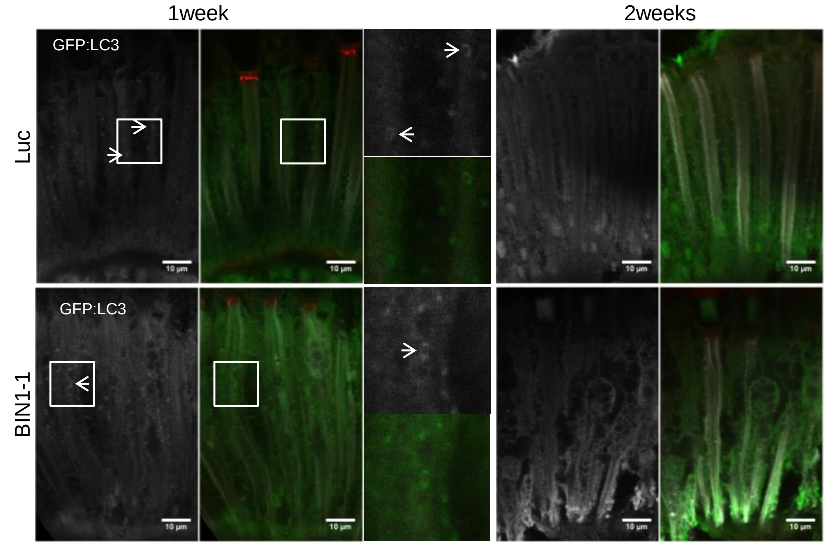

FIGURE S6: Percentage of cells in organoids

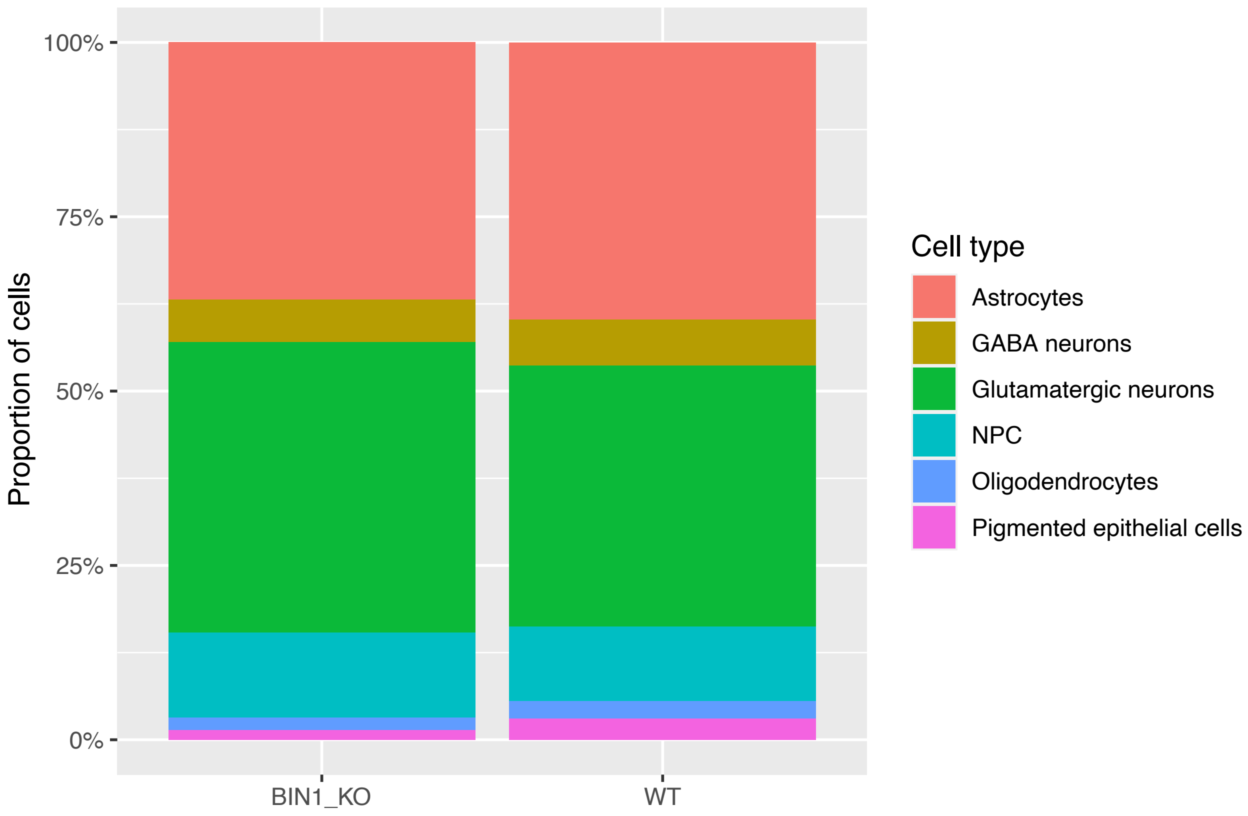
